# Modulation of Flagellar Rotation in Surface-Attached Bacteria: A Circuit for Rapid Surface-Sensing

**DOI:** 10.1101/567438

**Authors:** Maren Schniederberend, Jessica F. Johnston, Emilee Shine, Cong Shen, Ruchi Jain, Thierry Emonet, Barbara I. Kazmierczak

## Abstract

Attachment is a necessary first step in bacterial commitment to surface-associated behaviors that include colonization, biofilm formation, and host-directed virulence. The Gram-negative opportunistic pathogen *Pseudomonas aeruginosa* can initially attach to surfaces via its single polar flagellum. Although many bacteria quickly detach, some become irreversibly attached and express surface-associated structures, such as Type IV pili, and behaviors, including twitching motility and biofilm initiation. *P. aeruginosa* that lack the GTPase FlhF assemble a randomly placed flagellum that is motile; however, we observed that these mutant bacteria show defects in biofilm formation comparable to those seen for non-motile, aflagellate bacteria. This phenotype was associated with altered behavior of Δ*flhF* bacteria immediately following surface-attachment. Forward and reverse genetic screens led to the discovery that FlhF interacts with FimV to control flagellar rotation at a surface, and implicated cAMP signaling in this pathway. Although cAMP controls many transcriptional programs in *P. aeruginosa*, the known targets of this second messenger were not required to modulate flagellar rotation in surface-attached bacteria. Instead, alterations in switching behavior of the motor appear to result from previously undescribed effects of cAMP on switch complex proteins and/or the motor-stators associated with them.

**Author Summary:** Attachment to a surface often triggers programs of gene expression that alter the behavior, virulence and fitness of bacteria. Initial contact is usually mediated by surface exposed adhesins, such as flagella or pili/fimbriae, and there is much interest in how these structures might sense and respond to surface attachment. The human bacterial pathogen *Pseudomonas aeruginosa* usually contacts surfaces via its polar flagellum, the rotary motor that also powers bacterial swimming. We observed that wild-type bacteria quickly stopped rotating their flagellum after surface attachment, but that a mutant lacking the flagellar-associated protein FlhF did not. Using a combination of genetic approaches, we demonstrated that FlhF interacts with a component of the flagellar rotor (FliG) and with a polar scaffolding protein that positively regulates cAMP production (FimV) to stop flagellar rotation and thereby favor bacterial persistence at a surface. We provide evidence that the second messenger cAMP is the likely signal generated by flagellar-mediated surface attachment and show that cAMP is sufficient to alter the behavior of the flagellar motor.

## Introduction

Biofilms are clinically relevant, surface-associated multicellular bacterial communities. In the Gram-negative opportunistic pathogen *Pseudomonas aeruginosa*, a single polar flagellum and Type 4 pili (T4P) are essential for biofilm initiation and maturation, respectively [1]. Reversible, flagellar-mediated attachment of bacterial cells to a surface is somehow sensed, resulting in cessation of flagellar rotation and a transition to irreversible attachment [2, 3]. This step precedes surface-associated growth and T4P-mediated surface colonization, precursors to the formation of a matrix-encased biofilm community. The initial event of surface sensing is essential for this process, but how this bacterial sense of “touch” is perceived and transmitted remains unknown. A favored hypothesis is that increased flagellar load associated with surface tethering serves as a signal for surface attachment [4], but mediators that could transmit such a signal have not been definitively identified.

One of the earliest described examples of flagellar mechanosensing occurs in *Vibrio parahaemolyticus*. Lateral flagella (*laf*) genes, required for swarming motility, are transcribed when bacteria are transferred from liquid culture to high viscosity liquid or surface growth conditions [5]. Expression of a *laf::lux* transcriptional reporter can be induced by adding antibodies that specifically bind to and inhibit rotation of the organism’s polar flagellum, suggesting that motion of the polar flagellum may be sensed by Vibrio [6]. Fla-mutants, which fail to assemble a polar flagellum, exhibit constitutive, surface-independent *laf::lux* expression, consistent with the hypothesis that flagellar rotation/function—and perturbations thereof—can be sensed by a bacterium [6].

Examples of flagellar mechanosensing have since been provided in other systems, where they control not only switches between swimming and swarming behavior (e.g. *Proteus mirabilis* [7]), but also between swimming and adhesion (*e.g. Caulobacter crescentus* [8]) or biofilm initiation (*e.g. Bacillus subtilis* [9]). The ability of the flagellar motor to respond to alterations in load is clearly demonstrated by the load-dependent, step-wise recruitment of force-generating units to the *Escherichia coli* flagellar motor [4, 10, 11]. Thus most models of flagellar mechanosensing assume that alterations in flagellar load and function, as might occur in a high-viscosity environment or following flagellar-mediated attachment to a surface, initiate one or more potential signals such as altered proton flow across the motor, changes in stator conformation, or even cell envelope stress/deformation that are subsequently coupled to changes in cellular behavior [12].

The use of microfluidics to force bacterial proximity to a surface, coupled with real-time imaging of bacterial surface interactions, has allowed behaviors that bring bacteria to a surface to be separated from those required for surface sensing or surface adaptation [13, 14]. Thus pilus retraction, implicated in the *C. crescentus* transition from a motile “swarmer” cell to a surface-attached “stalked” cell [15], may contribute by bringing bacteria close to a surface, as pili are dispensable for rapid synthesis of the adhesive holdfast when cells are physically constrained near a surface [13]. In this setting, a component of the flagellar motor (MotB), but not the flagellar filament, hook or rod, is required for holdfast synthesis, and provides a signal that activates cyclic-di-GMP (cdGMP) synthesis [13]. Experiments that examine *P. aeruginosa* behavior immediately after surface attachment in a microfluidics device likewise implicate a cdGMP signal that is generated within seconds of attachment, in a process that depends on the MotAB flagellar motor-stator [14].

Although *P. aeruginosa* encodes only one flagellar system, it has two proton-driven motor-stators, MotCD (PA1460/PA1461) and MotAB (PA4954/PA4953), that can power its flagellum for swimming motility [2, 16–18] and play distinct roles in supporting swarming motility through media of increased viscosity [19]. Interactions of the MotAB and MotCD stators with the flagellar rotor are likely regulated, as a repressor of swarming motility, FlgZ, interacts with MotCD in a cyclic-di-GMP-dependent manner [20]. Regulators of flagellar placement (FlhF) and number (FleN/FlhG), while conserved among other polar flagellates [21], may also play unique roles linked to flagellar function in *P. aeruginosa* [22]. When *flhF*, which encodes a SRP-like GTPase, is deleted in *P. aeruginosa*, bacteria assemble a single flagellum that is no longer restricted to the pole [23]. Point mutants of FlhF that do not bind or hydrolyze GTP restore polar flagellar assembly in *P. aeruginosa* [22] and in *Shewanella oneidensis* [24], but affect flagellar motility in these organisms. In particular, *P. aeruginosa* expressing the GDP-locked FlhF(R251G) allele are paralyzed, despite assembling a polar flagellum [22]. This contrasts with phenotypes reported for *Campylobacter jejuni, Vibrio cholerae* and *V. alginolyticus*, where deletion of *flhF* results in predominantly aflagellate bacteria [25–28].

In this study we establish a role for FlhF in modulating flagellar behavior in surface-attached *P. aeruginosa*. We show that FlhF interacts with the polar organizer FimV and that this interaction is required for flagellar rotation to stop in surface-tethered bacteria. Using a combination of forward and reverse genetics to assess the roles of flagellar rotor and stator components in this process, we propose a model in which FlhF-FimV interactions are upstream of a cAMP-signaling pathway that alters flagellar function.

## Results

### FlhF is required to stop rotation of bacteria tethered at a surface

FlhF is a highly conserved signal recognition particle (SRP)-like GTPase important for placement and function of the *P. aeruginosa* polar flagellum [22, 23]. We made the unexpected observation that Δ*flhF* bacteria, which assemble a randomly placed but functional single flagellum, were as defective in biofilm formation as bacteria lacking a flagellum (Δ*fliC*) (Fig 1A). As the flagellum influences initial steps in biofilm formation, we developed a single-cell tethering assay to observe bacteria after surface attachment. The majority of tethered wild-type PAK bacteria (>80%) did not exhibit flagellar rotation after 2 minutes of incubation at an anti-flagellin antibody-coated surface (Fig 1B), and rotation stopped soon after 5 minutes in the remainder of the population (Fig 1C). In contrast, 58% of Δ*flhF* bacteria exhibited flagellar rotation at this initial 2 minute time-point, and the decay in the number of bacteria still rotating after 5 minutes was significantly slower than for wild-type organisms. Of note, flagellar rotation ceased with similar kinetics in wild-type bacteria and isogenic mutants lacking T4P (ΔpilA). As expected, we saw fewer Δ*pilA* bacteria detach during this assay, consistent with prior observations of *P. aeruginosa* behavior after flagellar-mediated surface attachment (Fig 1B & 1C). Thus wild-type *P. aeruginosa* quickly stop flagellar rotation after tethering, and Δ*flhF* is required for this behavior.

**Figure 1.**
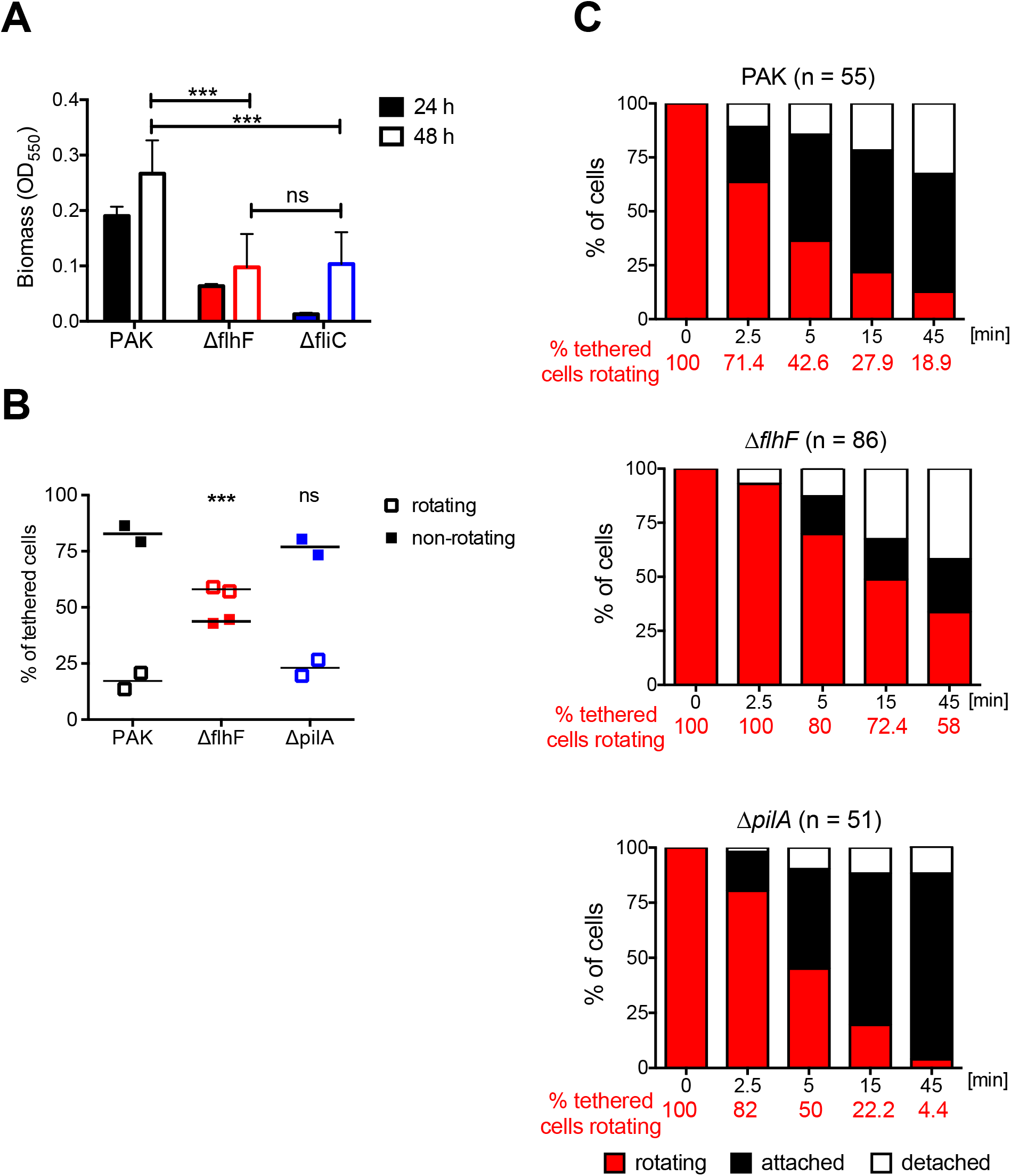
FlhF affects surface behaviors of *P. aeruginosa*. (A) Δ*flhF* cells are deficient in biofilm formation. Biomass of static biofilms was measured by crystal violet staining after 24 (solid) and 48 hours (open bars). Bars show mean ± S.D. (n = 4). Both Δ*flhF andΔfliC* differ from wild-type PAK (*** *p* < 0.001), but not from each other (2 way ANOVA with Bonferroni post-test). **(B) Δ*flhF* cells are more likely to rotate after surface tethering**. Percentage of rotating vs. non-rotating cells was determined 5 min after tethering to anti-FliC coated slides for PAK (n = 151), Δ*flhF* (n = 239) and Δ*pilA* (n = 135) bacteria. Each symbol represents an independent experiment; lines indicate means for each condition. The percentage of rotating Δ*flhF* cells, but not of Δ*pilA*, differs significantly from PAK (***, *p* < 0.001; 2 way ANOVA with Bonferroni post-test). **(C) Δ*flhF* rotation persists over time**. Rotating bacteria were identified 5 minutes after tethering to anti-FliC coated slides (t=0) and observed for 45 min. The proportion of rotating cells (red), attached cells (black), and detached cells (white) was determined in 4-8 independent experiments for PAK, Δ*flhF* and Δ*pilA*. Survival curves of rotating cells were analyzed with a Mantel-Cox test. The percentage of rotating Δ*flhF* cells differed from PAK or Δ*pilA* (***, *p* < 0.0001); the percentage of rotating PAK v Δ*pilA* cells did not differ significantly (*p* = 0.36).

### FlhF interacts with the C ring protein FliG

The flagellar base is formed by the inner membrane MS ring (composed of multiple copies of FliF), the C ring (FliG, FliM and FliN), and the export apparatus (reviewed in [29]). In the assembled flagellum, the C ring controls both torque generation (via interactions with motor/stator complex proteins) and the direction of flagellar rotation (via interactions with CheY-P). Since FlhF appeared to affect flagellar rotation, we used bacterial two-hybrid (B2H) to test whether FlhF interacted directly with components of the flagellar rotor or motor. Only FliG, the most membrane-proximal component of the C ring, interacted with FlhF in this assay (Fig 2). No significant variation in the strength of the B2H signal was observed when point mutant alleles of FlhF deficient in GTP binding or hydrolysis were tested for FliG binding (S1 Fig). Thus, FlhF localizes at the flagellar base [23] and appears to interact with the C ring protein FliG.

**Figure 2.**
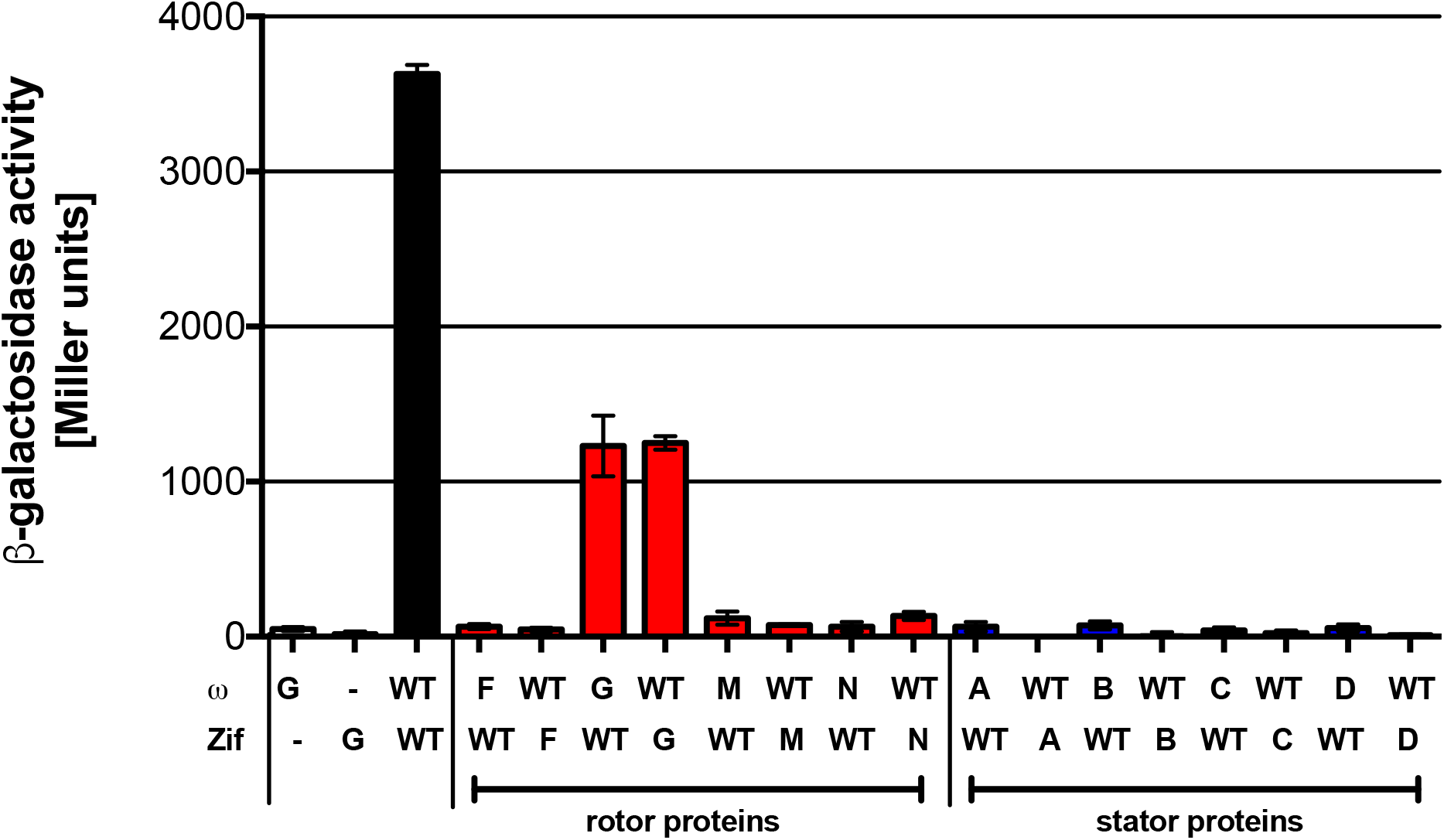
The C ring component FliG interacts with FlhF. FlhF interactions with flagellar proteins were assayed by bacterial two-hybrid assay. ω or Zif fusions were tested as indicated, with interactions resulting in beta-galactosidase expression and activity (reported in Miller units). Bars show mean ± S.D. (n =3) for a representative experiment. FlhF (WT), which forms a homodimer, served as a positive control (black bar). FliG (G) interacted with wild-type FlhF (WT) (red bars), but gave no signal when co-expressed with either the ω or Zif domain alone (white bars). Other tested rotor components (red bars: FliF (F), FliM (M), or FliN (N)) and motor-stator proteins (blue bars: MotA (A), MotB (B), MotC (C), or MotD (D)) did not show interactions with FlhF.

### FlhF interacts with FimV to stop flagellar rotation

The deletion of *flhF* resulted in bacteria that showed abnormally persistent flagellar rotation at a surface; however, we had previously observed that a mutation in the FlhF active site, of arginine 251 to glycine (R251G), results in loss of flagellar swimming motility [22]. FlhF(R251G) is “trapped” in a GDP-bound form [22] and has a dominant-negative (DN) effect on wild-type bacterial swimming motility (S2 Fig). We therefore hypothesized that FlhF might interact with other proteins to stop flagellar rotation, and that this interaction was absent in Δ*flhF* bacteria and inappropriately robust in bacteria expressing the DN allele. We tested this model by carrying out a screen for extragenic suppressors that would restore swimming motility to paralyzed, FlhF(R251G)-overexpressing bacteria. 23 independent suppressors were mapped by whole-genome sequencing to *vfr*, a cAMP-dependent transcriptional regulator, while 4 additional suppressors mapped to *fimV*, a protein whose expression is positively regulated by Vfr [30](S1 Table).

The Vfr mutations found in the suppressor strains either prematurely truncated this protein or altered residues implicated in Vfr dimerization, cAMP binding or DNA binding [31]. We tested these suppressors for their ability to twitch, a form of T4P-dependent motility that requires Vfr-dependent transcription of T4P structural and regulatory proteins, and found all to be twitching-negative, consistent with a loss of Vfr function (S3A Fig). Complementation *in trans* with an episomal copy of wild-type *vfr* both restored twitching motility (S3A Fig) and reverted the suppressor strains back to a paralyzed swimming phenotype (S3B Fig).

Because of the strong association between Vfr and twitching, we tested whether the loss of T4P was sufficient to suppress the paralyzed swimming phenotype associated with FlhF(R251G). This was not the case, however, as a mutant lacking the major pilin, *pilA*, was unable to swim when FlhF(R251G) was overexpressed (S4 Fig). Loss of Vfr function leads to an approximately two-fold increase in transcription of the master flagellar activator, FleQ [30], raising the possibility that overexpression of flagellar genes could suppress the paralyzed phenotype of FlhF(R251G) bacteria. However, steady-state FleQ protein levels were indistinguishable in parental and extragenic suppressor strains (S5A Fig), and increased expression of FleQ was not sufficient to restore swimming motility to FlhF(R251G) overexpressing bacteria (S5B Fig). Thus, the known effects of Vfr on expression of T4P and flagella do not account for the phenotype of the suppressor strains.

FimV, the second site of extragenic suppressor mutations, is also dependent on Vfr for its expression [30]. FimV is a large (919 aa) protein with an amino-terminal peptidoglycan binding domain, a single transmembrane helix, and a large cytoplasmic domain containing three tetratricopeptide repeats separated by an unstructured region between repeats 1/2 and 3 (Fig 3A) [32]. FimV is homologous to *Vibrio cholerae* HubP, a polar organizer [33]; in *P. aeruginosa* FimV positively regulates adenylate cyclase activity via an unknown mechanism and promotes normal T4P assembly and function [34]. We mapped one extragenic suppressor to a missense mutation (L7P) within the predicted signal sequence of FimV. This single point mutation was introduced into the endogenous *fimV* gene of wild-type *P. aeruginosa* by homologous recombination and found to be sufficient to suppress the paralyzed swimming phenotype associated with FlhF(R251G) overexpression (Fig 3B). We confirmed that *fimV(L7P)* bacteria had reduced twitching motility and decreased levels of intracellular cAMP, phenotypes described for *fimV* null mutants [34] and consistent with our observation that steady-state levels of FimV(L7P) were < 5% of the wild-type protein (S6 Fig). We also found that a loss-of-function transposon insertion mutant, PA14 *fimV*::Tn, was able to suppress the DN phenotype associated with FlhF(R251G) overexpression (Fig 3B). The suppressor phenotype was specific to *fimV*, as transposon insertions that disrupted the T4P assembly secretin (*pilQ*::Tn), or that disrupted components of the Pil/Chp chemotaxis cluster that can also increase intracellular cAMP in bacteria (*pilI*::Tn, *pilJ*::Tn, *pilG*::Tn) [34] were unable to suppress FlhF(R251G)-associated paralysis (Fig 3C). Thus, the loss of FimV specifically suppresses the DN swimming phenotype associated with FlhF(R251G) overexpression.

**Figure 3.**
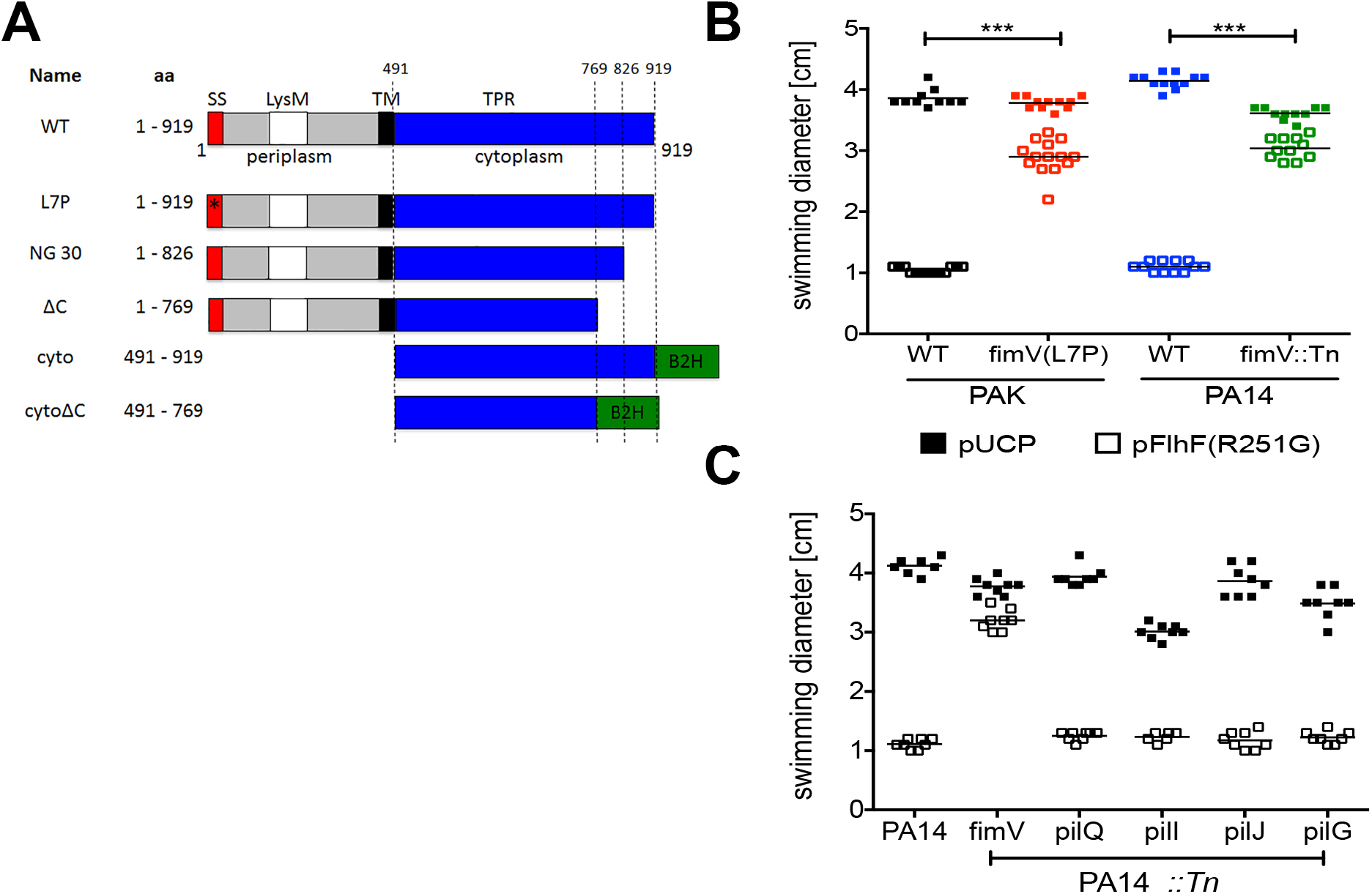
FimV(L7P) is a suppressor of FlhF(R251G). **(A) FimV constructs**. The predicted signal sequence (SS, red), LysM domain (white), periplasmic region (grey), transmembrane helix (TM, black) and tetratricopeptide repeats (TPR) (blue) are diagrammed. ω or Zif fusions for bacterial 2-hybrid screen are also shown (B2H, green). **(B) Loss of FimV function suppresses FlhF(R251G)**. Swimming zone diameters of PAK (black), PAK *fimV(L7P)* (red), PA14 (blue) and PA14 *fimV::Tn* (green) harboring either pUCP (vector control (VC)) or pFlhF(R251G) (open) are shown; each symbol is a biological replicate (line indicates mean). Disruption of *fimV* significantly alters swimming behavior of FlhF(R251G)-overexpressing bacteria (***, *p* < 0.001; 2 way ANOVA followed by Bonferroni posttest). **(C) Pil/Chp system mutants associated with low intracellular [cAMP] do not suppress FlhF(R251G)**. Wild-type and transposon mutant bacteria harboring harboring either empty vector (VC; solid) or pFlhF(R251G) (open) were assayed for swimming motility. Each symbol is a biological replicate; line indicates mean. Only disruption of *fimV* suppresses the dominant-negative phenotype associated with FlhF(R251G) overexpression (***, *p* < 0.001; 2 way ANOVA with Bonferroni posttest).

### FlhF interacts with the carboxy terminus of FimV

We had predicted that our suppressor screen would identify interaction partners of FlhF important for the regulation of flagellar rotation. We used B2H to test for interactions between FlhF, FimV and Vfr. We found no evidence of direct protein-protein interactions between FlhF and Vfr in this assay, but did demonstrate a robust interaction between FlhF and the cytoplasmic domain of FimV (aa 491-919) that was stronger when FlhF carried the R251G mutation (Fig 4A). One extragenic suppressor mapped to a nonsense mutation predicted to truncate FimV after aa 826, suggesting that the extreme carboxy-terminus of FimV, which contains its third tetratricopeptide repeat motif, might be required for FlhF-FimV interactions. Deletion of the final 150 aa of the FimV B2H construct (FimV aa 491-769) abolished FlhF-FimV interaction in the B2H assay (Fig 4A). We also used homologous recombination to replace *P. aeruginosa fimV* with the truncated *fimV*(Δ*C*) allele. This strain, PAK *fimV*(Δ*C*), was able to swim even when the DN FlhF(R251G) allele was overexpressed (S7 Fig). Together, these observations suggested that FlhF interactions with the carboxy-terminus of FimV lead to cessation of flagellar rotation.

**Figure 4.**
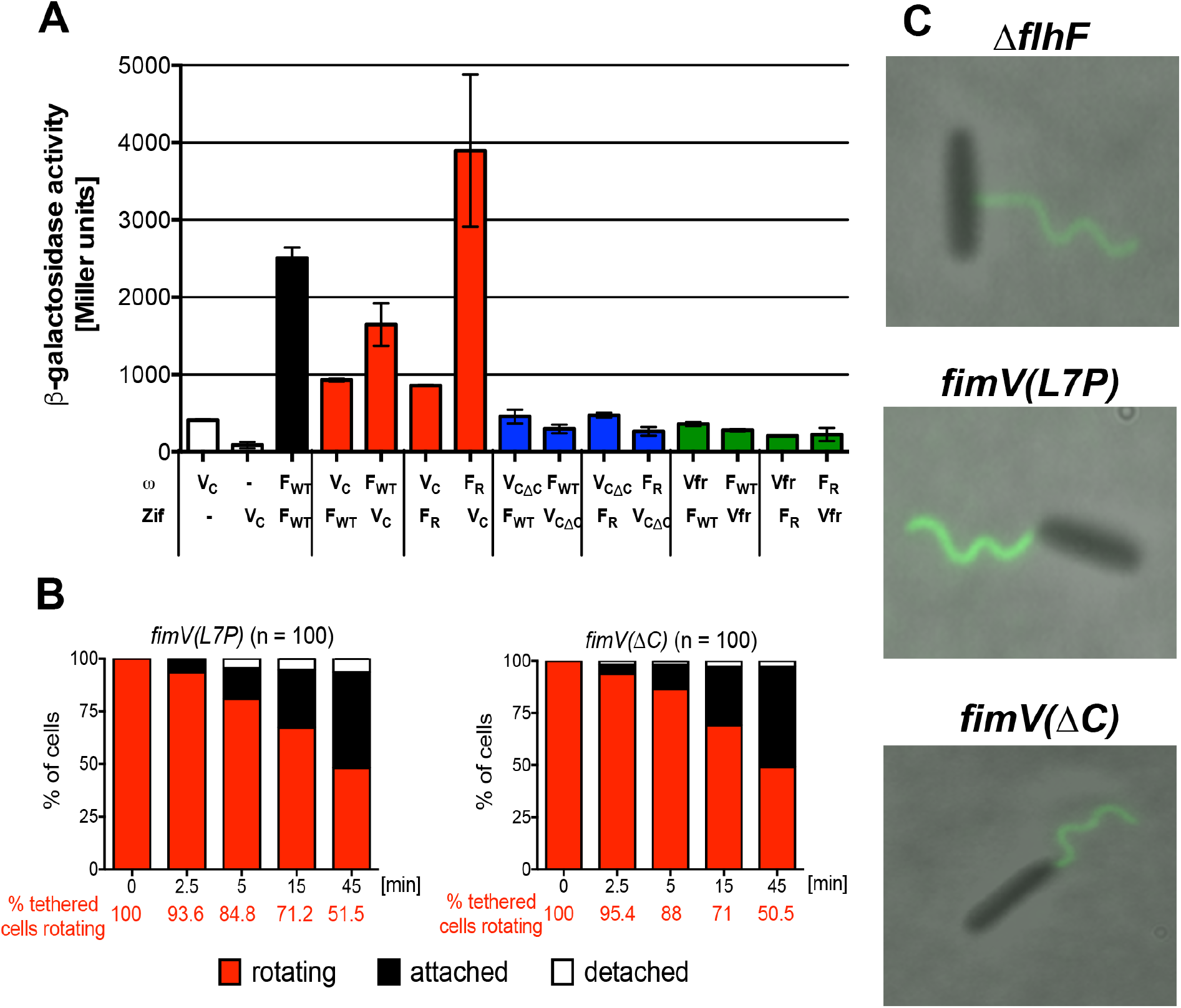
The carboxy terminus of FimV mediates interactions with FlhF and is required to stop rotation of tethered bacteria. **(A) FlhF interacts with the carboxy terminus of FimV**. FlhF interactions with FimV and Vfr were assayed by bacterial two-hybrid. ω or Zif fusions were constructed as indicated, with interactions resulting in beta-galactosidase expression and activity (reported in Miller units). Bars show mean ± S.D. (n =3) for a representative experiment. FlhF (F_WT_), which forms a homodimer, served as a positive control (black bar). FimV (V_C_) interacted with both wild-type (F_WT_) and R251G alleles (F_R_) of FlhF (red bars), but gave no signal when co-expressed with either the ω or Zif domain alone (white bars). Neither allele of FlhF interacted with a FimV construct lacking the final 150 aa (V_CδC_, blue bars) nor with Vfr (green bars). **(B) Surface-tethered bacteria expressing *fimV(L7P)* or *fimV*(Δ*C*) exhibit persistent rotation**. Rotating bacteria identified 5 minutes after tethering on anti-FliC coated slides (t=0) were observed for 45 min. The proportion of rotating cells (red), attached cells (black), and detached cells (white) was determined over 3-8 independent experiments. Survival curves of rotating *fimV(L7P)* and *fimV(ΔC*) differed significantly from PAK (*p* < 0.0001), but not from Δ*flhF* (p = 0.53 and p = 0.52, respectively; Mantel-Cox test). **(C) *fimV(L7P)* and *fimV(ΔC)* assemble a unipolar flagellum**. Cells were labelled with anti-FliC antibodies conjugated to Alexa Fluor 488 and visualized by phase and fluorescence microscopy.

### FimV is required to stop flagellar rotation in surface tethered bacteria

Surface-tethered Δ*flhF* bacteria are defective in stopping flagellar rotation. The hypothesis underlying our suppressor screen predicted that the protein that interacted with FlhF(R251G) to inhibit flagellar rotation inappropriately during swimming would also be required to stop flagellar rotation in surface-tethered cells. We therefore tested whether a *fimV* mutant phenocopied Δ*flhF* bacteria tethered at a surface, and found this to be the case for *fimV(L7P)* expressing cells (Fig 4B). Bacteria expressing the FimV allele lacking the carboxy-terminal domain required for FimV-FlhF interaction (PAK *fimV*(Δ*C*)) also showed persistent rotation after surface tethering (Fig 4B). In aggregate, these observations strongly supported the hypothesis that FlhF interacts with FimV to modulate flagellar rotation. Both *fimV(L7P)* and *fimV*(Δ*C*) bacteria assembled a polar flagellum, unlike Δ*flhF* bacteria, demonstrating that defective slowing of surface-tethered bacteria is not an artifact associated with assembly of a non-polar flagellum (Fig 4C).

### cAMP complements the persistent rotation phenotype of *flhF* and *fimV* mutants

FimV positively regulates the activity of the adenylate cyclase associated with T4P assembly and retraction in surface-associated *P. aeruginosa* [32, 35]. Addition of exogenous cAMP to *P. aeruginosa* adenylate cyclase mutants can complement phenotypes that depend on intracellular cAMP signaling [34]. We therefore tested whether extracellular cAMP would be sufficient to stop the rotation of Δ*flhF* or *fimV(L7P)* bacteria after surface tethering, and found this to be the case (Fig 5).

**Figure 5.**
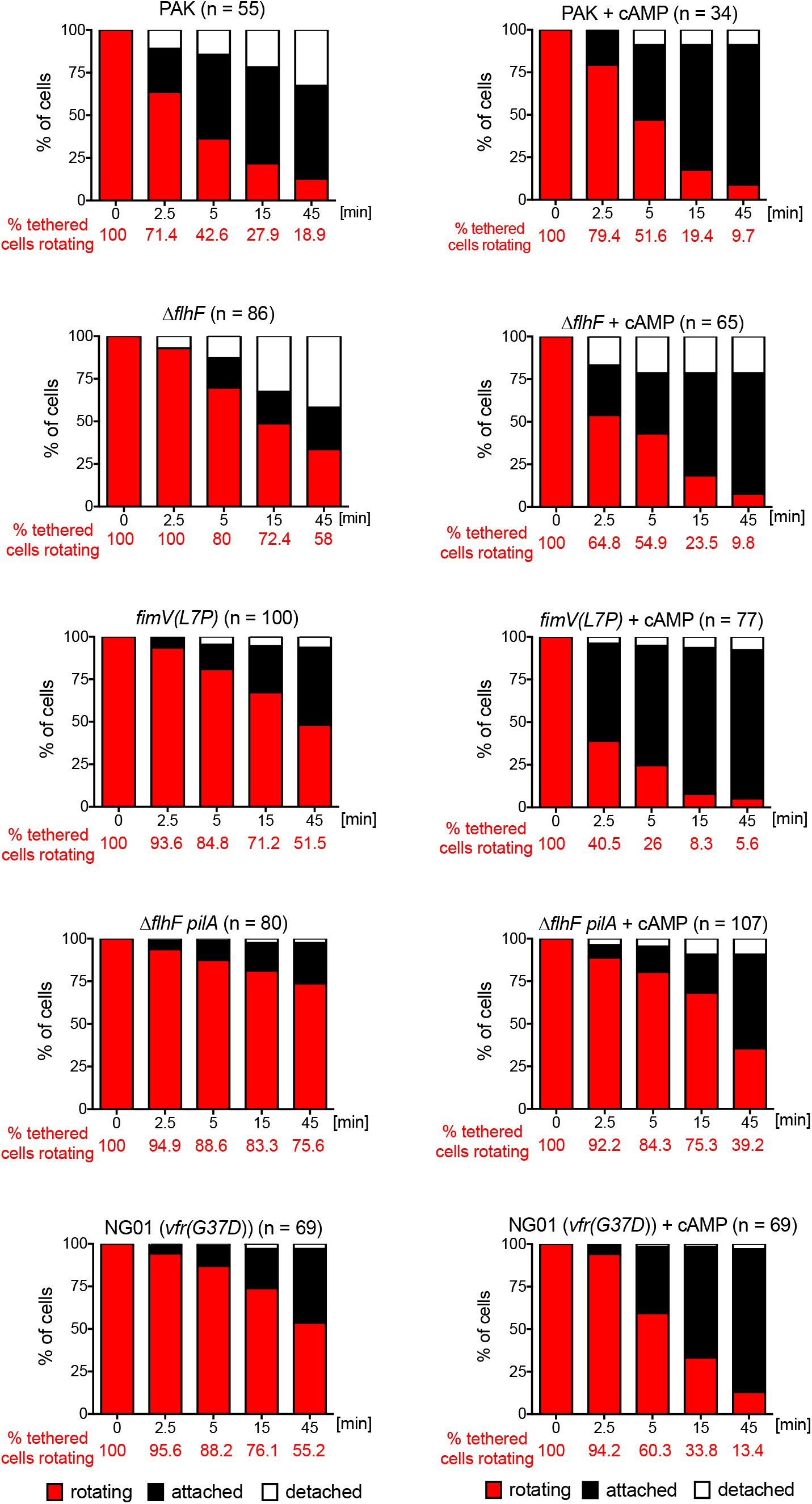
Exogenous cAMP complements the persistent rotation phenotype of surface-tethered *flhF, fimV* and *vfr* mutants. Bacteria were incubated with anti-FliC coated slides for 5 min in the presence or absence of 20 mM cAMP. Rotating bacteria were identified (t=0) and observed for 45 min. The proportion of rotating (red), attached (black), and detached cells (white) was determined in 3-8 independent experiments. Survival curves were analyzed with the Mantel-Cox test to determine whether exogenous cAMP had a significant effect on persistence of rotation. This was the case for Δ*flhF, fimV(L7P), NG01(vfr(G73D))* and Δ*flhF pilA* (*p* < 0.0001), but not for the parental strain PAK (*p* = 0.64).

The cAMP-dependent transcription factor Vfr had also been identified in our suppressor screen, although we did not find evidence of a direct interaction between it and FlhF (Fig 4). Nontheless, we confirmed that *vfr* mutants were also defective in stopping flagellar rotation after surface tethering (Fig 5). Surprisingly, exogenous cAMP could still complement this phenotype, similar to what we observed for *fimV* mutant bacteria. These observations suggested that protein(s) dependent on Vfr for their expression probably participate in regulating flagellar rotation at a surface, and that cAMP has a target other than Vfr in this process.

Polar pili have been implicated as just-in-time adhesins for *Caulobacter crescentus*, as their assembly by bacteria after surface binding stops flagellar rotation and plays a role in promoting permanent attachment of the organism [8, 15]. We tested whether T4P played an analogous role in *P. aeruginosa* surface attachment by comparing the behavior of surface bound Δ*flhF* and Δ*flhF pilA* organisms. In the absence of T4P, flagellar rotation persisted even longer, suggesting that pili contribute to stopping rotation of tethered bacteria. However, exogenous cAMP could still complement this defect, again arguing that cAMP is likely to have target(s) other than Vfr and T4P in this pathway (Fig 5).

### A mutation that increases flagellar load raises intracellular cAMP levels in bacteria

The ability of cAMP to modulate the behavior of tethered Δ*flhF* and *fimV(L7P)* bacteria led us to test whether increased flagellar load, which bacteria experience when tethered to a surface by their flagellum, is associated with increased intracellular cAMP. To generate a uniformly high load in all bacteria, we deleted *fleN*, a regulator of flagellar number in *P. aeruginosa* and other polar flagellates [36]. As expected, these bacteria assembled thick bundles of polar flagella (Fig 6A). Intracellular cAMP levels as measured by ELISA were increased in Δ*fleN* bacteria as compared to wild-type PAK (Fig 6B). This increase in cAMP was not observed in the absence of FimV (Δ*fleN fimV(L7P)*), or when bacteria carried the Δ*fleN* mutation but could not assemble flagella (Δ*fleN* Δ*flhA*). Thus, endogenous cAMP levels increase in a FimV-dependent manner in bacteria experiencing high flagellar load.

**Figure 6.**
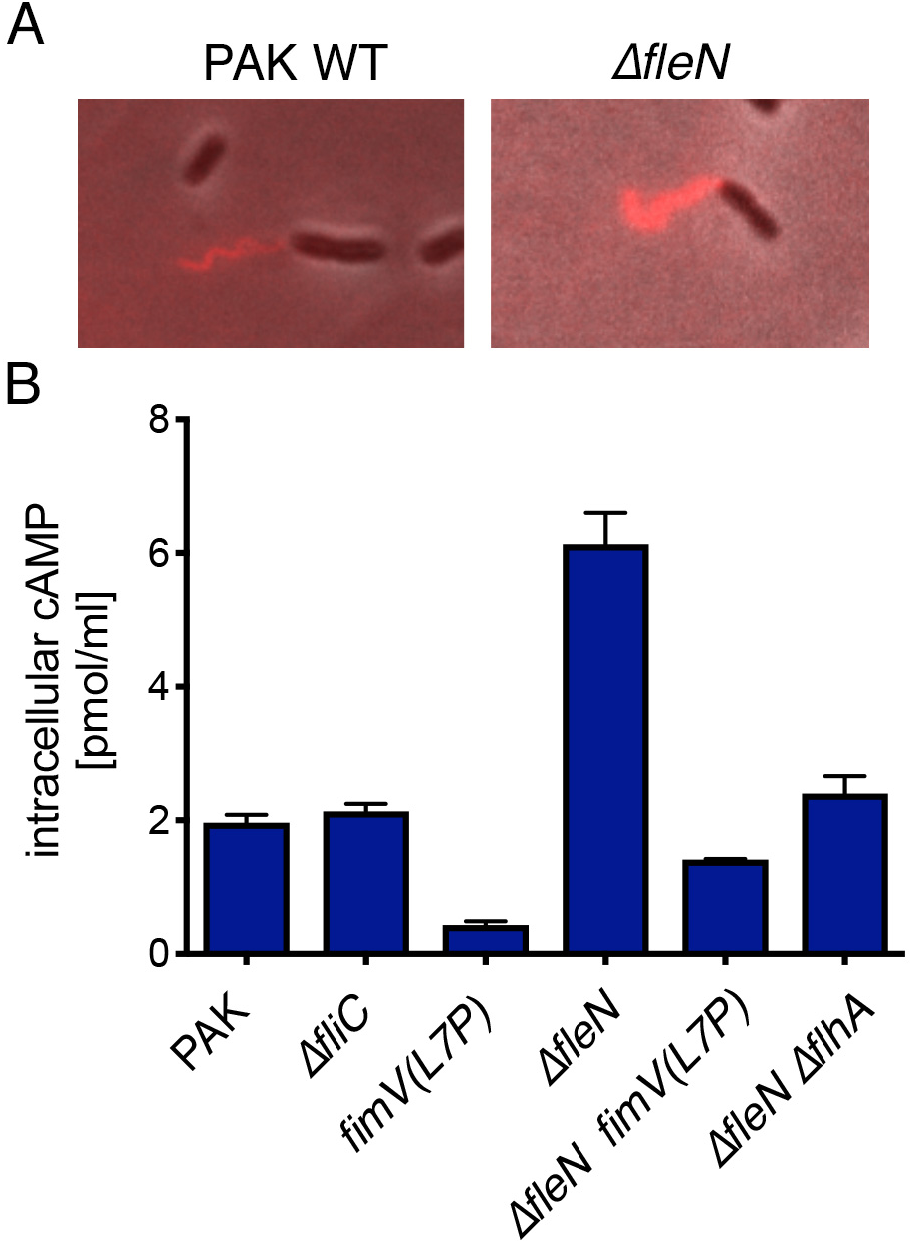
Increased flagellar load in Δ*fleN* bacteria increases cAMP in a FimV-dependent manner. **(A) ΔfleN bacteria assemble increased numbers of polar flagella**. Cells were labelled with anti-FliC antibodies conjugated to Alex Fluor 594 and visualized by phase and fluorescence microscopy. **(B) cAMP levels measured by ELISA**. Intracellular cAMP levels were measured by ELISA. Bars show mean ± s.d. of triplicate samples from a representative experiment of 3-5 independent assays.

### The MotCD motor is necessary for flagellar rotation in tethered bacteria, but the MotAB motor is required for rotation to stop

When swimming bacteria approach a surface, attach via their flagellum, and continue to spin, the flagellum experiences a significant increase in load. In *E. coli*, the flagellum responds to increased load by recruiting additional motor-stator pairs to the flagellum [4, 37]. *P. aeruginosa* is unusual in encoding two motor-stator pairs that interact with the polar flagellum to drive PMF-dependent motility, MotCD (PA1460/PA1461) and MotAB (PA4954/PA4953) [17, 18]. The two motor-stator pairs are not functionally equivalent, as only MotCD can support swarming motility [19]. We hypothesized that MotAB and MotCD might have distinct functions in swimming versus surface-tethered bacteria, and constructed strains lacking *motAB* or *motCD* to test this. We confirmed that our mutants displayed previously described phenotypes in plate-based swimming assays, where both are defective *vis-a-vis* wild-type bacteria, (S8 Fig) and in swarming assays (S9 Fig). Despite their similar phenotypes in swimming agar assays, the Δ*motAB* and Δ*motCD* bacteria exhibited markedly different swimming speeds and reversal behavior during swimming in liquid media. Speed and reversal frequency of the Δ*motCD* mutant, which expresses only MotAB, were close to indistinguishable from those of wild-type PAK, suggesting that swimming may be largely powered by the MotAB motor-stator in liquid (Fig 7). The Δ*motAB* mutant had a lower reversal frequency than either wild-type or Δ*motCD* bacteria, similar to that of a Δ*cheR1* methyltransferase mutant [38] that we constructed as a control, suggesting that the motor-stators influence chemotactic properties of the flagellum.

**Figure 7.**
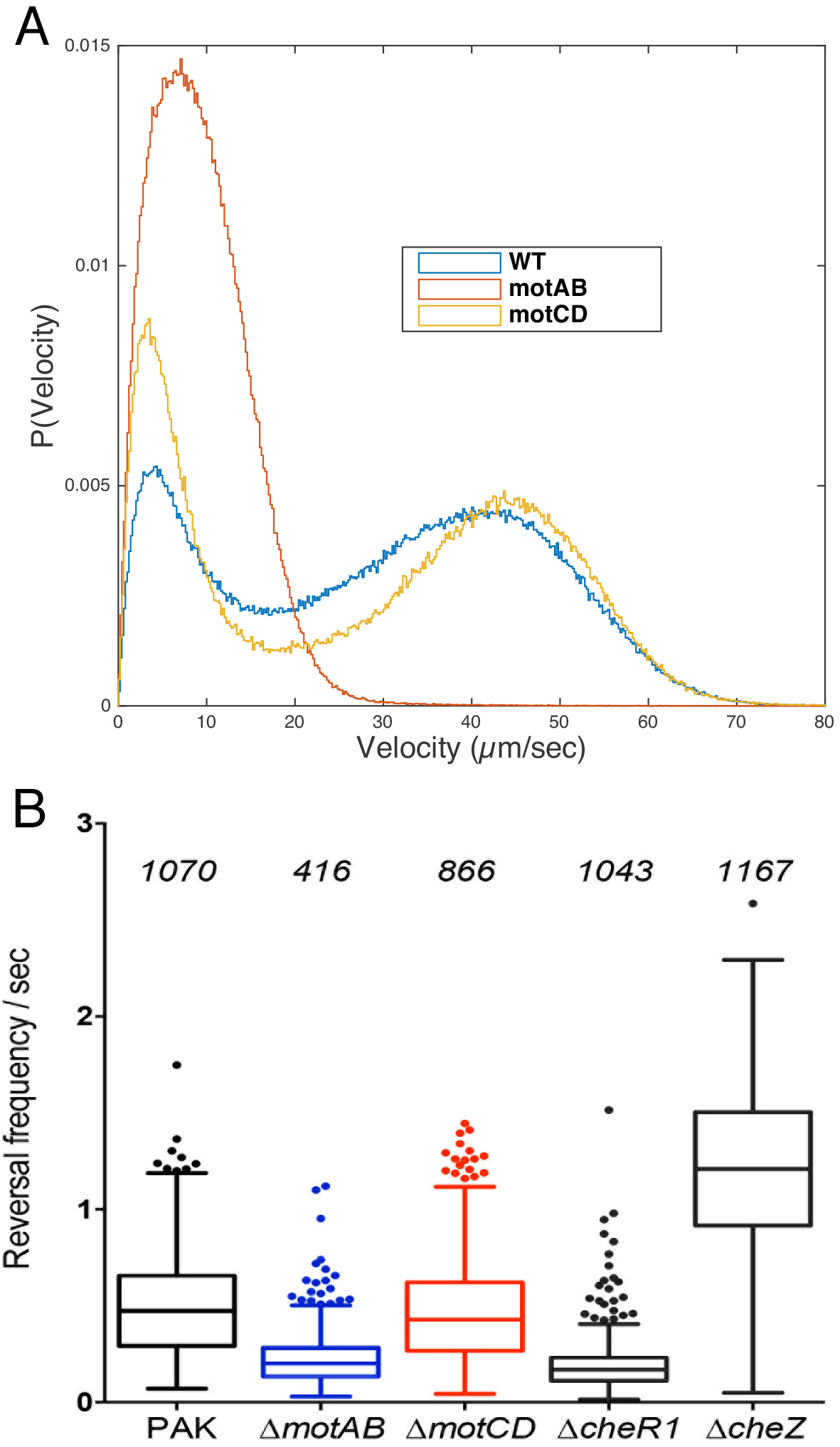
Δ*motAB* and Δ*motCD* bacteria show distinct distributions of velocity (A) and reversal frequency (B) during liquid swimming. Cells of the each genotype were imaged under dark-field microscopy at 30 fps. Image analysis was carried out as described in Methods using a custom MATLAB code.

We hypothesized that bacteria expressing only one of the two *P. aeruginosa* motor-stators might also differ in their behavior at a surface. Δ*motCD* bacteria could be tethered by anti-flagellin antibodies, but very few were able to rotate (S10 Fig), an observation consistent with the proposed role of MotCD in flagellar rotation under conditions of high external load [19]. Surprisingly, tethered Δ*motAB* bacteria exhibited prolonged flagellar rotation after tethering, despite the presence of FlhF and FimV (Fig 8A). We found that the addition of exogenous cAMP had a modest effect on the rotation of Δ*motAB* bacteria and no effect at all on the rotation of Δ*flhF motAB* bacteria. These findings demonstrate that the MotCD motor-stator is required for tethered bacteria to spin, but that the MotAB motor is necessary for rotation to stop (Fig 8A).

**Figure 8.**
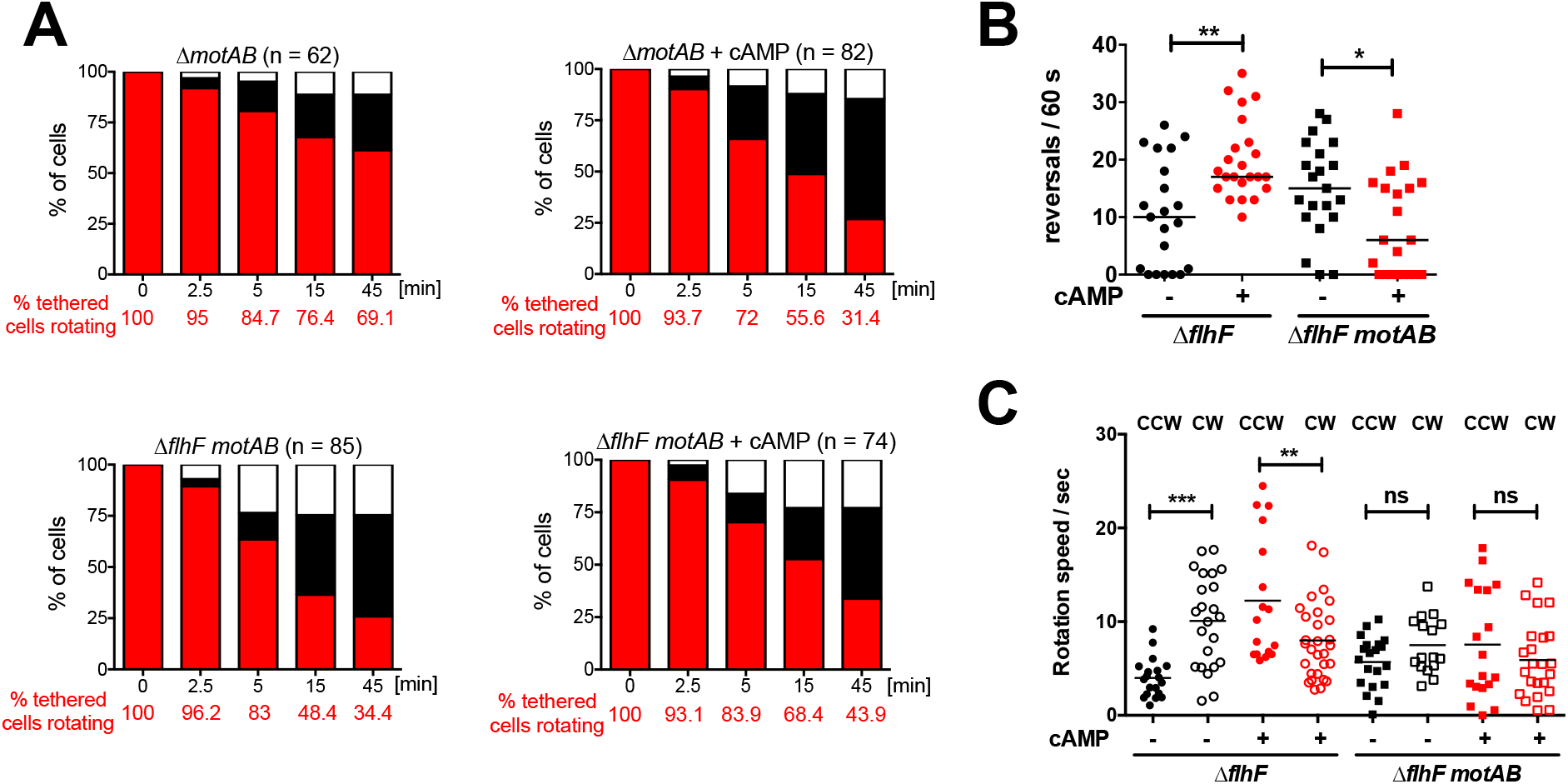
The MotAB stator is required for tethered bacteria to stop rotating. Bacteria were incubated with anti-flagellin antibody coated slides for 5 min in the presence or absence of 20 mM cAMP. **(A) Behavior of tethered bacteria over time**. Videomicroscopy was used to identify bacteria rotating at t=0, and the behavior of these cells was followed for 45 min. The proportion of rotating cells (red), attached cells (black), and detached cells (white) was determined in 4-8 independent experiments. To compare the percentage of tethered cells that were rotating in presence or absence of cAMP, survival curves were analyzed with a Mantel-Cox test. Exogenous cAMP had a significant effect on PAK Δ*motAB* (***, *p* = 0.0005), but not on PAK Δ*flhF* Δ*motAB* (ns, *p* = 0.0916). **(B) Reversal frequency of tethered bacteria**. Movies of tethered bacteria imaged at 100 fps for 60s were analyzed by a custom Matlab script to determine reversal frequency. Each symbol indicates reversals made by an individual cell over 60s (n = 21-23); lines indicate medians. cAMP increases the reversal frequency of Δ*flhF*, but decreases reversal frequency of the Δ*flhF motAB* mutant (Kruskal-Wallis with Dunn’s multiple comparison test; *, *p* <0.05; **, *p* < 0.01). **(C) Rotation speeds of tethered bacteria**. CW and CCW rotation speeds were extracted from tethered bacteria analyzed in panel (B).

### cAMP changes flagellar reversal behavior in tethered bacteria

We used high-speed video-microscopy to examine more closely the behavior of tethered Δ*flhF* and Δ*flhF motAB* bacteria before and after addition of exogenous cAMP. cAMP addition changed the behavior of the Δ*flhF* rotor in several ways. The median number of reversals observed in tethered bacteria increased significantly (Fig 8B), and CCW rotation speed increased significantly (Fig 8C). In contrast, when tethered Δ*flhF motAB* were exposed to cAMP, no change in median reversal frequency or rotation speed was observed. Thus observed changes in rotor behavior of tethered bacteria in response to cAMP depend on the presence of the MotAB motor-stator. In aggregate, these findings suggest that the MotAB motor might itself be a direct or indirect target of cAMP.

## Discussion

Bacterial binding to a surface via the flagellum is a first, necessary step in the transition to surface-associated behaviors such as T4P mediated surface colonization and biofilm formation [2, 16]. Ultimately, bacteria commit to these surface-associated behaviors by downregulating flagellar gene expression and upregulating the expression of adhesins and biofilm-matrix components [3]. However, modulation of flagellar rotation, rather than flagellar production, is a strategy that allows bacteria first to reversibly bind to and sample a surface [39–42].

We have demonstrated that FlhF, a GTPase required for polar flagellar placement, is also necessary for flagellar rotation to stop after surface binding. Using unbiased genetic screens we found that the cessation of flagellar rotation required the HubP-like protein FimV, which interacted with FlhF (Fig 4). An interaction between FlhF and HubP has previously been reported in *V. cholerae*, although neither protein affects the other’s distribution and ~95% of Δ*hubP* bacteria assemble a single polar flagellum[33]. In *P. aeruginosa*, FimV is a well-known positive regulator of adenylate cyclase activity [34], and we found that exogenous cAMP could rescue the defect of either Δ*flhF* or *fimV* mutant bacteria in stopping flagellar rotation after surface-tethering. cAMP is a pleiotropic regulator that interacts with the transcription activator Vfr to upregulate T4P gene expression and also positively regulates T4P-mediated motility, via proteins of the Chp/Pil chemotaxis-like cluster, as recently reviewed [43]. Thus, we were surprised to observe that cAMP still stopped flagellar rotation in bacteria lacking Vfr or pili. These findings suggested that cAMP has targets other than T4P or Vfr in this pathway, while subsequent experiments demonstrated that cAMP alters flagellar reversal frequency and rotation speed in a MotAB dependent manner (Figs 7, 8).

*P. aeruginosa* is unusual in encoding two motor-stator pairs that both interact with the polar flagellum to drive PMF-dependent motility [17, 18]; in other bacteria, different motor-stators may associate with polar vs. lateral flagella, or drive flagellar rotation in response to proton vs. Na+ gradients [29]. Prior work has suggested that the two motor-stators of *P. aeruginosa* are recruited to the flagellar rotor under distinct conditions [19], and our observations agree with this hypothesis. Wild-type and Δ*motCD* bacteria swimming through a low viscosity medium have very similar swimming speeds and reversal frequencies, in contrast to Δ*motAB* cells, suggesting that the MotAB motor primarily powers swimming under this condition (Fig 7). However, when bacteria were tethered to a surface via their flagellum, continued rotation in this condition of increased drag required the presence of MotCD (S11 Fig). Unexpectedly, surface-tethered bacteria were less likely to stop rotating if they lacked MotAB, and exogenous cAMP could not complement this phenotype. High-speed tracking of tethered cells showed that cAMP increased reversal frequency of the flagellum through a mechanism that depended on MotAB (Fig 8). Although we have yet to identify the direct target of cAMP in this pathway, our data in aggregate suggest that cAMP may result in MotAB recruitment to the rotor of tethered cells, thereby altering its behavior.

Mechanisms to change flagellar behavior have been reported for many different bacteria, and often coincide with bacterial adaptation to surface-attached growth [39–42, 44]. These changes in flagellar behavior are often mediated by cyclic-di-GMP (cdG) effectors that interact with the motor-switch complex [45]. Many mechanisms increase cdG levels in bacteria. These are often associated with activation of diguanylate cyclases in response to bacterial surface contact [46], and can occur quite rapidly, as has recently been reported following bacterial surface attachment in microfluidics devices [14]. The behavior that we have characterized precedes association of *P. aeruginosa* bacteria with a surface and appears to increase the probability that surface attachment will occur. Our findings are consistent with the recent observation that MotAB is required for rapid polar localization of FimW, an event that reflects a rapid rise in intracellular cdG in surface-attached bateria [14], as we observe that MotAB is necessary for bacteria to stop flagellar attachment after tethering and become surface-attached.

We propose that swimming bacteria that bind to a surface via their flagellum experience increased load on the flagellar motor. *P. aeruginosa* tethered to a surface can continue rotating, but only if they express the MotCD motor stator, suggesting that this motor-stator is recruited to the flagellum under this condition. Wild-type bacteria quickly stop flagellar rotation and become surface-attached, a process that requires interaction between FlhF and FimV. Our data suggest that a cAMP signal is produced as a result of FlhF-FimV interaction, and that cAMP modulates reversal frequency and speed of the flagellar motor in a MotAB-dependent manner. We speculate that MotAB may be recruited back to the flagellum in this setting, leading to the observed changes in flagellar behavior which may help drive the bacteria toward the surface [47]. These events likely precede the cdG-dependent signaling cascade described by Jenal and colleagues, which requires MotAB for its initiation [14].

The roles of second-messenger signaling in *P. aeruginosa* surface-association are complex and likely highly redundant. cAMP signals in surface-associated bacteria are amplified by the Pil/Chp system, which mechano-senses via the retraction of T4P [48, 49], and lead to surface-associated motility and virulence. Transient “surface-exposure” by swimming cells, however, also changes cAMP levels in individual bacteria that fail to commit to a surface, as recently demonstrated by Lee *et al*. [50]. These so-called “surface-sentient” cells exhibit higher levels of intracellular cAMP and are more likely to progress to surface association and irreversible attachment on subsequent encounters with a surface. Whether this cAMP-dependent “memory” requires the FlhF-FimV circuit that we have described remains to be tested.

## Materials and Methods

### Bacterial strains, media and culture conditions

Strains and plasmids used in this study are listed in S1 Table. Bacteria were propagated in Luria Broth (LB) (1% tryptone, 0.5% yeast extract, 1% NaCl), on LB agar, or on Vogel-Bonner minimal medium (VBM) agar plates [51]. Antibiotics were added to liquid and solid media as appropriate at the following concentrations: *E. coli*, 100 μg/ml ampicillin, 15 μg/ml gentamicin, 50 μg/ml kanamycin, and 20 μg/ml tetracycline; *P. aeruginosa*, 200 μg/ml carbenicillin, 100 μg/ml gentamicin, and 100 μg/ml tetracycline.

### Static biofilm formation

Static biofilm assays were performed as previously described [52]. Briefly, overnight cultures grown in LB were diluted into fresh medium (1:100); 100 μl of the dilution was used to inoculate 3 to 4 replicates per strain in a 96 well plate (Costar #2797). Plates were incubated for 24 h at 30°C, then gently washed to remove non-adherent bacteria. Adherent biofilm was stained with crystal violet (0.1%), followed by biofilm dissolution with glacial acetic acid (1%). The amount of dissolved crystal violet was measured as absorbance at 550 nm. Each experiment was repeated independently at least 3 times.

### Extragenic suppressor screen

An exponential phase (OD_600_ = 0.4) culture of *P. aeruginosa* strain PAK Δ*flhF attB::flhF(R251G)* was divided and treated with either 0.1 M ethyl methane sulfonate (EMS), 0.1 mg/ml methylnitronitrosoguanidine (NG), or irradiated with UV light (1-4 times with 100,000 Joules). One aliquot was left untreated to allow for emergence of spontaneous mutations. After further incubation at 37°C for 30, 60, 90, or 120 minutes, cells were sampled, pelleted, washed twice, and then resuspended in LB. Each bacterial pool was frozen in 15% (v/v) glycerol at −80°C.

To screen for suppressors of the paralyzed phenotype of the parental strain, aliquots of mutagenized cells were inoculated in LB medium overnight and then spotted on 0.3% LB swimming agar plus 2% arabinose. Swimming “plates” were cast in sterile reagent reservoirs (Costar #4870), allowing bacteria and serine chemoattractant to be spotted at opposite ends of the reservoir. After incubating swimming plates overnight at 30°C, bacteria were sampled at 0.5 cm intervals from the origin by stabbing agar with a 200μl sterile pipet tip and resuspending the plug in 50 μl of LB. Each of these samples was plated onto LB agar, and the one furthest from the origin that still showed bacterial growth was used to inoculate swimming plates for a second round of selection. After this second swimming round, bacteria were again sampled at different distances from the origin and frozen as glycerol stocks at −80°C. Pools were subsequently streaked for single colonies and approximately 200 single colonies were subjected to further analysis.

Intragenic *flhF(R251G)* suppressors were excluded by transforming each candidate with a plasmid overexpressing FlhF(R251G). Only candidates that were still able to swim were further evaluated. Twenty of the strongest suppressors were subjected to whole genome sequencing to map the sites of suppressor mutations, along with the parental strain PAK Δ*flhF attB::flhF(R251G)* and wild-type PAK.

### Whole genome sequencing and sequence analysis

Whole genome sequencing was carried out as previously described [53]. Briefly, bacterial genomic DNA was prepared for sequencing on the Illumina MiSeq by the Yale Center for Genomic Analysis using the TruSeq DNA LT Sample Prep Kit (Illumina). Purified libraries were barcoded, pooled, and sequenced on the MiSeq using a 2 × 250 paired-end protocol. Initial basecalls were converted to fastq files using MiSeq CASAVA software suite, demultiplexed, and clipped for adaptors. Sequences were examined using FastQC (Galaxy) [54]. Reads were mapped to the *P. aeruginosa* PAO1 reference genome with the BWA (v. 0.6.2) software package using default parameters [55, 56]. Bowtie (v. 2.1.0) was used to align reads [57], and SNPs were called using the Samtools mpileup [58].

### Construction of chromosomal mutations

Mapped mutations were introduced into the chromosome of the parental strain, PAK Δ*flhF attB::flhF(R251G)*, or of wild-type PAK (as indicated) by homologous recombination. Briefly, primers flanking the mutation of interest were synthesized to contain *attB1* or *attB2* sites compatible with the Gateway Clonase II (Invitrogen) system. The mutation plus a linked, “silent” loss or gain of a restriction site was engineered into overlapping internal primers, and these four primers were used to generate overlapping PCR products that were spliced together by overlap extension PCR (S2 Table). PCR products were cloned into the Gateway-adapted suicide vector pDONRX [34], screened by restriction mapping, and confirmed by DNA sequencing. Constructs containing only the desired mutation were transformed into *E. coli* S17-1 and mobilized into *P. aeruginosa* by mating.

Exconjugants were selected on VBM gentamicin and then streaked to VBM plus 10% sucrose to select for loss of vector backbone sequences through a second recombination event [59]. Sucrose-resistant, gentamicin-susceptible colonies were screened by amplifying the targeted gene and digesting the PCR product to look for the linked gain/loss of a restriction site. Colonies that passed this screen were sequenced to confirm the presence of the desired mutation in the endogenous chromosomally encoded gene. This strategy was also used to introduce a BB2 epitope tag prior to the stop codon of the chromosomally encoded *fimV* and *fimV(L7P)* genes.

The PAK Δ*flhF pilA* strain was constructed by allelic recombination, using a previously described pEX18Gm *pilA::aacC1* suicide vector in which the *pilA* gene is replaced by a gentamicin resistance cassette [60]. Briefly, this construct was mobilized into PAK Δ*flhF* by mating. Exconjugants were selected on VBM-gentamicin plus 10% (w/v) sucrose (to select for gene replacement and loss of the vector-carried *sacB* gene). The mutation was confirmed by PCR, Western blotting, and phenotype (loss of twitching motility).

Gene deletions of *motA/motB, motC/motD, cheR1*, and *cheZ* were introduced into the chromosome by homologous recombination. Briefly, primers flanking the deletion of the gene of interest were synthesized to contain *attB1* or *attB2* sites compatible with the Gateway Clonase II (Invitrogen) system. PCR products were cloned into the Gateway-adapted suicide vector pDONRX [34]. The procedures for generating ex-conjugants, screening candidates, and confirming the presence of the desired mutation(s) are described above.

### Western blotting

Western blotting was carried out as previously described [23]. Membranes were probed with antisera against FleQ, FlhF, Hfq, and BB2-epitope tag (1:4,000) followed by HRP-conjugated goat anti-rabbit antibody (1:4,000; Bio-Rad). Membrane blocking, washes, and visualization of bound antibody by enhanced chemiluminescence were carried out as previously described [23]. Signals were detected using an Image Station 4000R (Kodak) and quantified with Carestream Molecular Imaging software (version 5.0.2.28).

### Motility assays

Swimming assays were performed by spotting 5 μl of a fresh overnight culture of bacteria (grown at 37°C with aeration in LB) diluted to OD_600_ = 1 onto 0.3% LB agar plates supplemented with antibiotics or 20mM cAMP when appropriate. The diameter of the swimming zones was measured after overnight incubation (16 h) at 30°C.

For swarming assays, M8 minimal medium plates (0.5% agar) supplemented with 0.4% glucose and 0.05% sodium glutamate were used. Swarming plates were inoculated with 5 μl of an overnight culture (diluted as for swimming assays), incubated at 30°C overnight (16 h), and then incubated at room temperature for an additional 24 h. Pictures of the whole agar plates were taken for comparison with an Image Station 4000R (Kodak).

Twitching assays were performed by stabbing a colony through 1% LB agar to the agar/plastic interface with a sterile toothpick. The diameters of twitching zones were measured after 24 hours of incubation at 37°C, at which time the twitching zone was visible on the plastic surface. All motility assays were performed at least three times using 4 or more replicates per experiment.

### Bacterial two-hybrid assay

Genes of interest were subcloned into pBRGPω and pACTR-AP-Zif to generate carboxy terminal fusions, respectively, with the ω subunit of *E. coli* RNA polymerase or the Zn-finger DNA binding domain of murine Zif268 [61]. FlhF wild-type and mutant alleles were cloned into both bacterial two-hybrid vectors as previously described (Schniederberend et al., 2013). The cytoplasmic domain of FimV (FimV_C_ or V_C_, amino acids 491-919) was PCR amplified from PAK genomic DNA and cloned the same way using the corresponding primers NdeI-FimV-forw and NotI-FimV-rev (Table S2). A truncated version of this FimV domain lacking the C-terminal region (FimV_ΔC_ or V_CΔC_, amino acids 491-769) was cloned using the reverse primer NotI-FimV2-rev. Full-length Vfr was cloned using the primers NdeI-Vfr-forw and NotI-Vfr-rev. Flagellar genes of interest (*fliF, fliG, fliM, fliN, motA, motB, motC*, and *motD*) were PCR amplified from PAK genomic DNA and cloned the same way into both bacterial two-hybrid vectors using the corresponding primers NdeI-gene-forw and NotI-gene-rev. All constructs were confirmed by DNA sequencing.

Plasmids containing ω- and Zif fusions were electroporated simultaneously into *E. coli* KΔZif1ΔZ, which carries a *lacZ* reporter gene downstream of a promoter containing the Zif binding site [61]. Expression of the fusion proteins was induced by 50 μM IPTG. β-galactosidase activity was measured as described previously [62] and repored in Miller units. Samples were assayed in triplicate in three independent experiments.

### Fluorescence microscopy

Flagella were stained as previously described [22]. Briefly, *P. aeruginosa* overnight cultures grown at 37°C in LB were diluted 1:100 into fresh medium the next morning, and grown with aeration for 2h at 37°C. Cells were harvested by centrifugation (3,000 *g* for 3 min), fixed in 4% paraformaldehyde for 20 min at room temperature and washed twice with phosphate-buffered saline (PBS). Flagella were stained with anti-FliC antibodies labeled with Alexa Fluor 488 (1 μg/ml in PBS) for 30 min at room temperature, then washed with PBS. Stained flagella were visualized using a Nikon Eclipse TS100 microscope (100x objective) equipped with a fluorescein isothiocyanate filter and a monochrome Spot camera (Diagnostic Instruments) running Spot 4.0.1 software.

### FliC tethering assay

FliC binding assays were performed as previously described [22]. Briefly, *P. aeruginosa* was grown overnight at 37°C in LB medium, diluted 1:50 into fresh medium the next morning, and incubated for 1h at 37°C. Slides were incubated with PBS containing BSA (100 μg/ml) plus anti-FliC antibodies (2 μg/ml) for 30 min at RT. Antibody-coated slides were incubated with bacteria (and with 20 mM cAMP, as indicated) for 5 min at RT prior to imaging.

For the analysis of the cell fate over time, movies were taken immediately using a Nikon Eclipse TS100 microscope (100x objective) equipped with a Canon Vixia HFS200 camera. To monitor the same cells over time, 30 sec movies were taken at 2-15 minute intervals. All cells rotating at t=0 were tracked in each subsequent movie and scored for one of three fates: attached/rotating, attached/not rotating, detached.

### Swimming analysis

Bacteria were grown overnight at 37°C in LB, diluted 1:100 into fresh medium the next morning, and incubated for 2h at 37°C. Cells were spotted onto glass slides and observed by dark-field microscopy with a Zeiss Axiostar plus microscope (10x objective). Video clips were obtained using a Canon Vixia HFS200 camera. For each strain, the movement of at least 100 bacteria was analyzed for 1 min (1800 consecutive images with 30 frames per 1 sec) to obtain data for the swimming analysis.

To identify objects, the movies were background-subtracted by averaging over six-second windows and subtracting that average from each frame in that window. To analyze individual cell speed and angular velocity, each cell’s trajectory was reconstructed from the movie image sequence using a custom MATLAB (Mathworks) code [63]. This tracking software uses particle detection via a radial symmetry method [64] and particle tracking adapted from u-track 2.1 [65]. The cell speed at each time point was calculated according to

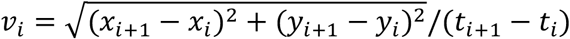

and the cell angular velocity was calculated according to

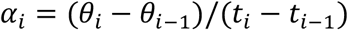

normalizing for the frame rate of the camera. To remove non-cell tracks, we filtered the tracks to have a mean speed between 15 and 80 μm/s (except for Δ*motAB*, where a mean speed filter of 10-80 μm/s was used), a ≥ 3 sec trajectory time, and a maximum mean squared displacement of ≥ 400 μm^2^.

#### Reversal analysis, swimming cells

To define reversals in free-swimming bacteria, a modification of the algorithm described by Theves *et al*. was written in MATLAB [66]. All changes in angular velocity

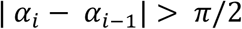

were identified and considered potential reversals, and the maximum cell speed measured between all potential reversals was extracted. For a potential reversal x to be scored a true reversal, the following criteria all needed to be met: (i) the maximum cell speed measured during the interval between (x-1, x) was >20 μm/s (15 μm/s for Δ*motAB*); (ii) the maximum cell speed measured during the interval between (x, x+1) was >20 μm/s (15 μm/s for Δ*motAB*); (iii) the minimum speed within 0.1s of the potential reversal x was less than half of the maximum cell speed during the interval before or after the reversal; (iv) no other potential reversals were identified within 0.1s before or after x. These parameters filtered out “pausing” or “jiggling” events, where bacteria exhibit repeated large changes in angular velocity without moving [66]. Speed, angular velocity, and true reversals were plotted for each track as a function of time and manually examined to confirm expected behavior of our algorithm.

To calculate reversal frequency, the number of true reversals per track was divided by the track length. In cases of zero reversals, we calculated an upper limit for reversal frequency by assuming that one reversal would be observed if the tracking time was extended by 0.3s. The percentage of Δ*motAB* tracks affected by this correction is 10.7-11.3%; for Δ*cheR1*, it is 39-46% (with vs. without exogenous cAMP).

### Reversal frequency analysis, tethered cells

For the analysis of reversal frequency, bacteria and anti-FliC coated slides were prepared as described above. Tethered cells were recorded for 60 seconds at 100 frames per second with a digital scientific CMOS camera (Hamamatsu ORCA-Flash4.0 V2, 2×2 pixel binning, 1024 × 1024 array, 10 ms exposure) mounted on an inverted microscope (Nikon Eclipse TI-U) with a 100x oil immersion objective (Nikon CFI Plan Fluor, N.A. 1.30, W.D 0.2 mm) and LED white light diascopic illumination (Thorburn Illumination Systems).

Movies of tethered cells were analyzed frame by frame using a custom MATLAB (Mathworks) script, which fits an ellipse over the cell body, measures the length of the ellipse’s major axis, and calculates the centroid (center of mass) of the ellipse. These values were then used to obtain the cell body’s angle (in degrees) with respect to the center of the image region. This step was repeated for each subsequent frame of the movie. Once all angles were known for each movie frame, the angular velocity and rotation frequency was calculated by

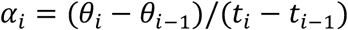

and

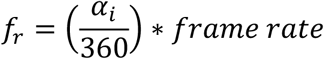

and reversal frequency was obtained from the switching frequency from clockwise to counterclockwise rotation (and vice-versa).

### Statistical analysis

Curve fitting and data analysis were carried out using Prism 5.0 (GraphPad) software. Data are expressed as means ± SD. For data sets in which two conditions were varied, *p* values were calculated by analysis of variance (two way ANOVA) followed by Bonferroni’s posttest. Normally distributed data sets were compared by one-way ANOVA followed by Bonferroni’s posttest, while data sets that were not normally distributed were analyzed by Kruskal-Wallis and Dunn’s Multiple Comparison posttest. For statistical analysis of survival curves the Mantel-Cox test was performed. *p* values < 0.05 were considered significant.

## Acknowledgments

We thank Ruben Ramphal for anti-FleQ antiserum, and George O’Toole for PA14 transposon mutants used in this study. We thank members of the Kazmierczak lab for thoughtful discussion and comments.

## Supporting Information

**Figure S1. FliG-FlhF interactions are observed with FlhF catalytic site mutants**. ω or Zif fusions to FliG and to wild-type and mutant alleles of FlhF were constructed as indicated, with interactions resulting in beta-galactosidase expression and activity (reported in Miller units). Bars show mean ± S.D. (n =3) for a representative experiment. The FlhF homodimer (“WT”), serves as a positive control (black bar). FliG (“G”) interacted with all tested alleles of FlhF, including the hydrolytically active wild-type (“WT”) and FlhF(L298R,P299L) (“LP”) alleles, the GDP-locked FlhF(R251G) (“R”), as well as alleles defective in GTP hydrolysis (FlhF(K222A), “K”) or binding (FlhF(D294A), “D”). No signal was observed when FliG was co-expressed with either the ω or Zif domain alone (white bars).

**Figure S2. FlhF(R251G) has a dominant negative effect on swimming**. A second copy of *flhF* or *flhF(R251G)* was integrated into the *attB* site of PAK and expressed from an inducible arabinose promoter. Swimming zone diameter was determined in the presence of 0.2% (open symbols) and 0.4% arabinose (solid symbols); lines indicate means for each condition. Overexpression of FlhF(R251G) significantly inhibited swimming motility (***, *p* < 0.001; 2way ANOVA with Bonferroni post-test).

**Figure S3. Motility analysis of suppressors mapped to *vfr***. Suppressor mutants mapped to *vfr* were transformed with plasmid-encoded wild-type Vfr (red) or empty vector (black) and assayed for motility. Missense mutations and the amino acid position preceding indels are indicated for each suppressor. **(A) Twitching motility of suppressors is complemented *in trans* by wild-type Vfr**. Each symbol represents the median of 6-10 technical replicates; the error bar shows the interquartile range. Complementation with wild type Vfr had a significant effect on twitching motility of all suppressor mutants, but not on the parental strain PAK Δ*flhF* + *attB::flhF(R251G)* (ns, *p* > 0.05). **(B) Complementation of *vfr* suppressors *in trans* with wild-type Vfr reverts cells to a paralyzed swimming phenotype**. Each symbol shows median ± interquartile range of ≥ 10 technical replicates. Over-expression of wild type Vfr had a significant effect on all suppressor strains, but not on the parental strain. (Two-way ANOVA with Bonferroni post-test; *, *p* < 0.05; **, *p* < 0.01; ***, *p* < 0.001; *n.s.,p* > 0.05.)

**Figure S4. Absence of Type IV pili does not suppress the FlhF(R251G) phenotype**.

FlhF(R251G) was over-expressed in wild-type PAK and the isogenic *pilA* mutant. In both strains the dominant negative effect of FlhF(R251G) on swimming was observed. Each point represents a technical replicate swimming assay; lines indicate means.

**Figure S5. FleQ levels are unchanged in suppressors and have no effect on swimming motility**. (A) Lysates prepared from overnight cultures grown in LB + 2% arabinose (ca. 1 × 10^7^ cells/lane) were separated by SDS-PAGE, transferred to PVDF and probed with antisera against FleQ, FlhF and Hfq (loading control). Chemiluminescence was used to detect and quantify antibody binding; the graph shows mean intensity ± SD for 3-6 replicates relative to the parent strain (parent (“P”), Δ*flhF* + *attB::FlhF(R251G))*. (B) Swimming motility was assayed on semisolid agar for wild-type and FlhF(R251G) overexpressing bacteria transformed with a FleQ expression construct (pUCP/FleQ) or vector control (pUCP). Each symbol indicates a replicate; the line indicates the mean.

**Figure S6. PAK *fimV(L7P)* destabilizes FimV.** The BB2 epitope tag was recombined at the carboxy-terminal end of the endogenous *fimV* gene or the *fimV(L7P)* allele as described in Methods. Whole cell lysates corresponding to 2 × 10^8^ cells (expect for lane 4, 6 × 10^8^ cells) were separated by SDS-PAGE, transferred to PVDF, and probed with anti-BB2 monoclonal antibody. Lane 1: PAK *fimV-BB2*, Lane 2: PAK, Lane 3: empty, Lane 4: PAK *fimV(L7P)-BB2*.

**Figure S7. Deletion of the carboxy terminal region of FimV suppresses the dominant negative swimming defect associated with FlhF(R251G)**. The carboxy-terminal domain of the chromosomal *fimV* gene was deleted, resulting in PAK *fimV*(Δ*C*). Overexpression of FlhF(R251G) from a plasmid inhibits swimming of PAK, but the dominant negative phenotype is significantly suppressed in the *fimV*(Δ*C*) background. Lines indicate the mean of 8-12 independent replicates. (***, *p* > 0.001; two-way ANOVA with Bonferroni posttest).

**Figure S8. Δ*motAB* and Δ*motCD* bacteria show defects in plate-based swimming assays**. *motAB* and *motCD* genes were deleted by homologous recombination, and swimming behavior of the resulting mutant strains was assayed on 0.3% LB agar. Swimming diameters were measured for mutants (carrying empty pUCP vector) and for complemented strains as indicated (n=6-8).

**Figure S9. MotCD is required for swarming in both wild-type and Δ*flhF* bacteria**. Unmarked deletions of the *motAB* and *motCD* genes were constructed by homologous recombination in PAK and Δ*flhF* backgrounds. Swarming was assayed on 0.5% agar as described in Methods.

**Figure S10. Δ*motCD* cells are able to attach to a surface, but do not spin**. Tethered cells were imaged by videomicroscopy 5 minutes after incubation on anti-flagellin antibody coated or BSA coated slides. Percentage of unattached (open symbol) vs. tethered (solid symbol) cells are shown for PAK, Δ*flhF*, Δ*motCD*, and Δ*flhF motCD* (n= 300-500 cells per strain/condition). The inset table shows the number of tethered vs. spinning cells for each strain with anti-flagellin coated slides.

**S1 Table. Mutations mapped in FlhF(R251G) suppressors**

**S2 Table. Bacterial strains and plasmids used in this study**

**S3 Table. Primers used in this study**

## References

1. O’Toole GA, Kolter R. Flagellar and twitching motility are necessary for *Pseudomonas aeruginosa* biofilm development. Mol Microbiol. 1998;30:295–304.

2. Conrad JC, Gibiansky ML, Jin F, Gordon VD, Motto DA, Mathewson MA, et al. Flagella and pili-mediated near-surface single-cell motility mechanisms in *P. aeruginosa*. Biophys J. 2011;100(7):1608–16. Epub 2011/04/06. doi: 10.1016/j.bpj.2011.02.020. PubMed PMID: 21463573; PubMed Central PMCID: PMC3072661.

3. Caiazza NC, O’Toole GA. SadB is required for the transition from reversible to irreversible attachment during biofilm formation by *Pseudomonas aeruginosa* PA14. J Bacteriol. 2004;186:4476–85.

4. Lele PP, Hosu BG, Berg HC. Dynamics of mechanosensing in the bacterial flagellar motor. Proc Natl Acad Sci U S A. 2013;110:11839–44.

5. Belas R, Simon M, Silverman M. Regulation of lateral flagella gene transcription in *Vibrio parahaemolyticus*. J Bacteriol. 1986;167:210–8.

6. McCarter L, Hilmen M, Silverman M. Flagellar dynamometer controls swarmer cell differentiation of V. parahaemolyticus. Cell. 1988;54:345–51.

7. Belas R, Suvanasuthi R. The ability of *Proteus mirabilis* to sense surfaces and regulate virulence gene expression involves FliL, a flagellar basal body protein. J Bacteriol. 2005;187(19):6789–803. doi: 10.1128/JB.187.19.6789-6803.2005. PubMed PMID: 16166542; PubMed Central PMCID: PMCPMC1251568.

8. Li G, Brown PJ, Tang JX, Xu J, Quardokus EM, Fuqua C, et al. Surface contact stimulates the just-in-time deployment of bacterial adhesins. Mol Microbiol. 2012;83(1):41–51. Epub 2011/11/08. doi: 10.1111/j.1365-2958.2011.07909.x. PubMed PMID: 22053824; PubMed Central PMCID: PMC3245333.

9. Cairns LS, Marlow VL, Bissett E, Ostrowski A, Stanley-Wall NR. A mechanical signal transmitted by the flagellum controls signalling in *Bacillus subtilis*. Mol Microbiol. 2013;90(1):6–21. Epub 2013/07/31. doi: 10.1111/mmi.12342. PubMed PMID: 23888912; PubMed Central PMCID: PMC3963450.

10. Chawla R, Ford KM, Lele PP. Torque, but not FliL, regulates mechanosensitive flagellar motor-function. Scientific reports. 2017;7(1):5565. doi: 10.1038/s41598-017-05521-8. PubMed PMID: 28717192; PubMed Central PMCID: PMCPMC5514156.

11. Tipping MJ, Delalez NJ, Lim R, Berry RM, Armitage JP. Load-dependent assembly of the bacterial flagellar motor. mBio. 2013;4(4). Epub 2013/08/22. doi: 10.1128/mBio.00551-13. PubMed PMID: 23963182; PubMed Central PMCID: PMC3747592.

12. Belas R. Biofilms, flagella, and mechanosensing of surfaces by bacteria. Trends Microbiol. 2014;22(9):517–27. Epub 2014/06/05. doi: 10.1016/j.tim.2014.05.002. PubMed PMID: 24894628.

13. Hug I, Deshpande S, Sprecher KS, Pfohl T, Jenal U. Second messenger-mediated tactile response by a bacterial rotary motor. Science. 2017;358(6362):531–4. Epub 2017/10/28. doi: 10.1126/science.aan5353. PubMed PMID: 29074777.

14. Laventie B-J, Sangermani M, Estermann F, Manfredi P, Planes R, Hug I, et al. A Surface-Induced Asymmetric Program Promotes Tissue Colonization by *Pseudomonas aeruginosa*. Cell Host & Microbe. 2019;25(1):140–52.e6. doi: https://doi.org/10.1016/j.chom.2018.11.008.

15. Ellison CK, Kan J, Dillard RS, Kysela DT, Ducret A, Berne C, et al. Obstruction of pilus retraction stimulates bacterial surface sensing. Science. 2017;358(6362):535–8. Epub 2017/10/28. doi: 10.1126/science.aan5706. PubMed PMID: 29074778; PubMed Central PMCID: PMCPMC5805138.

16. Toutain CM, Caizza NC, Zegans ME, O’Toole GA. Roles for flagellar stators in biofilm formation by *Pseudomonas aeruginosa*. Res Microbiol. 2007;158(5):471–7. doi: 10.1016/j.resmic.2007.04.001. PubMed PMID: 17533122.

17. Toutain CM, Zegans ME, O’Toole GA. Evidence for two flagellar stators and their role in the motility of *Pseudomonas aeruginosa*. J Bacteriol. 2005;187:771–7.

18. Doyle TB, Hawkins AC, McCarter LL. The complex flagellar torque generator of *Pseudomonas aeruginosa*. J Bacteriol. 2004;186:6341–50.

19. Kuchma SL, Delalez NJ, Filkins LM, Snavely EA, Armitage JP, O’Toole GA. Cyclic di-GMP-mediated repression of swarming motility by *Pseudomonas aeruginosa* PA14 requires the MotAB stator. J Bacteriol. 2015;197(3):420–30. doi: 10.1128/JB.02130-14. PubMed PMID: 25349157; PubMed Central PMCID: PMCPMC4285984.

20. Baker AE, Diepold A, Kuchma SL, Scott JE, Ha DG, Orazi G, et al. PilZ Domain Protein FlgZ Mediates Cyclic Di-GMP-Dependent Swarming Motility Control in *Pseudomonas aeruginosa*. J Bacteriol. 2016;198(13):1837–46. doi: 10.1128/JB.00196-16. PubMed PMID: 27114465; PubMed Central PMCID: PMCPMC4907108.

21. Kazmierczak BI, Hendrixson DR. Spatial and numerical regulation of flagellar biosynthesis in polarly flagellated bacteria. Mol Microbiol. 2013;88(4):655–63. Epub 2013/04/23. doi: 10.1111/mmi.12221. PubMed PMID: 23600726; PubMed Central PMCID: PMC3654036.

22. Schniederberend M, Abdurachim K, Murray TS, Kazmierczak BI. The GTPase Activity of FlhF Is Dispensable for Flagellar Localization, but Not Motility, in *Pseudomonas aeruginosa*. J Bacteriol. 2013;195(5):1051–60. Epub 2012/12/25. doi: 10.1128/JB.02013-12. PubMed PMID: 23264582.

23. Murray TS, Kazmierczak BI. FlhF is required for swimming and swarming in *Pseudomonas aeruginosa*. J Bacteriol. 2006;188(19):6995–7004.

24. Gao T, Shi M, Ju L, Gao H. Investigation into FlhFG reveals distinct features of FlhF in regulating flagellum polarity in *Shewanella oneidensis*. Mol Microbiol. 2015;98:571–85.

25. Correa NE, Peng F, Klose KE. Roles of the regulatory proteins FlhF and FlhG in the *Vibrio cholerae* flagellar transcription hierarchy. J Bacteriol. 2005;187:6324–32.

26. Balaban M, Joslin SN, Hendrixson DR. FlhF and its GTPase activity are required for distinct processes in flagellar gene regulation and biosynthesis in *Campylobacter jejuni*. J Bacteriol. 2009;191:6602–11.

27. Green JCD, Kahramanoglou C, Rahman A, Pender AMC, Charbonnel N, Fraser GM. Recruitment of the earliest component of the bacterial flagellum to the old cell division pole by a membrane-associated signal recognition particle family GTP-binding protein. J Mol Biol. 2009;391:679–90.

28. Kusumoto A, Nishioka N, Kojima S, Homma M. Mutational analysis of the GTP-binding motif of FlhF which regulates the number and placement of the polar flagellum in *Vibrio alginolyticus*. J Biochem. 2009;146(5):643–50. Epub 2009/07/17. doi: 10.1093/jb/mvp109. PubMed PMID: 19605463.

29. Minamino T, Imada K. The bacterial flagellar motor and its structural diversity. Trends Microbiol. 2015;23(5):267–74. doi: 10.1016/j.tim.2014.12.011. PubMed PMID: 25613993.

30. Wolfgang MC, Lee VT, Gilmore ME, Lory S. Coordinate regulation of bacterial genes by a novel adenylate cyclase signaling pathway. Devlop Cell. 2003;4:253–63.

31. Serate J, Roberts GP, Berg O, Youn H. Ligand Responses of Vfr, the Virulence Factor Regulator from *Pseudomonas aeruginosa*. J Bacteriol. 2011;193(18):4859–68. Epub 2011/07/19. doi: 10.1128/JB.00352-11. PubMed PMID: 21764924.

32. Buensuceso RN, Nguyen Y, Zhang K, Daniel-Ivad M, Sugiman-Marangos SN, Fleetwood AD, et al. The Conserved Tetratricopeptide Repeat-Containing C-Terminal Domain of *Pseudomonas aeruginosa* FimV Is Required for Its Cyclic AMP-Dependent and -Independent Functions. J Bacteriol. 2016;198(16):2263–74. Epub 2016/06/15. doi: 10.1128/jb.00322-16. PubMed PMID: 27297880; PubMed Central PMCID: PMCPMC4966435.

33. Yamaichi Y, Bruckner R, Ringgaard S, Moll A, Cameron DE, Briegel A, et al. A multidomain hub anchors the chromosome segregation and chemotactic machinery to the bacterial pole. Genes Dev. 2012;26(20):2348–60. Epub 2012/10/17. doi: 10.1101/gad.199869.112. PubMed PMID: 23070816; PubMed Central PMCID: PMC3475806.

34. Fulcher NB, Holliday PM, Klem E, Cann MJ, Wolfgang MC. The *Pseudomonas aeruginosa* Chp chemosensory system regulates intracellular cAMP levels by modulating adenylate cyclase activity. Mol Microbiol. 2010;76(4):889–904. Epub 2010/03/30. doi: 10.1111/j.1365-2958.2010.07135.x. PubMed PMID: 20345659; PubMed Central PMCID: PMC2906755.

35. Inclan YF, Persat A, Greninger A, Von Dollen J, Johnson J, Krogan N, et al. A scaffold protein connects type IV pili with the Chp chemosensory system to mediate activation of virulence signaling in *Pseudomonas aeruginosa*. Mol Microbiol. 2016;101(4):590–605. Epub 2016/05/05. doi: 10.1111/mmi.13410. PubMed PMID: 27145134; PubMed Central PMCID: PMCPMC4980298.

36. Dasgupta N, Arora SK, Ramphal R. *fleN*, a Gene That Regulates Flagellar Number in *Pseudomonas aeruginosa*. J Bacteriol. 2000;182(2):357–64. doi: 10.1128/jb.182.2.357-364.2000.

37. Tipping MJ, Delalez NJ, Lim R, Berry RM, Armitage JP. Load-dependent assembly of the bacterial flagellar motor. mBio. 2013;4:e00551–13. doi: 10.1128/mBio.00551-.

38. Schmidt J, Musken M, Becker T, Magnowska Z, Bertinetti D, Moller S, et al. The *Pseudomonas aeruginosa* chemotaxis methyltransferase CheR1 impacts on bacterial surface sampling. PLoS One. 2011;6(3):e18184. Epub 2011/03/30. doi: 10.1371/journal.pone.0018184. PubMed PMID: 21445368; PubMed Central PMCID: PMC3062574.

39. Boehm A, Kaiser M, Li H, Spangler C, Kasper CA, Ackermann M, et al. Second messenger-mediated adjustment of bacterial swimming velocity. Cell. 2010;141:107–16.

40. Paul K, Nieto V, Carlquist WC, Blair DF, Harshey RM. The c-di-GMP binding protein YcgR controls flagellar motor direction and speed to affect chemotaxis by a “backstop brake” mechanism. Mol Cell. 2010;38:128–39.

41. Blair KM, Turner L, Winkelman JT, Berg HC, Kearns DB. A molecular clutch disables flagella in the Bacillus subtilis biofilm. Science. 2008;320(5883):1636–8. Epub 2008/06/21. doi: 10.1126/science.1157877. PubMed PMID: 18566286.

42. Fang X, Gomelsky M. A post-translational, c-di-GMP-dependent mechanism regulating flagellar motility. Mol Microbiol. 2010;76(5):1295–305. doi: 10.1111/j.1365-2958.2010.07179.x. PubMed PMID: 20444091.

43. Leighton TL, Buensuceso RN, Howell PL, Burrows LL. Biogenesis of *Pseudomonas aeruginosa* type IV pili and regulation of their function. Environ Microbiol. 2015. Epub 2015/03/27. doi: 10.1111/1462-2920.12849. PubMed PMID: 25808785.

44. Guttenplan SB, Blair KM, Kearns DB. The EpsE flagellar clutch is bifunctional and synergizes with EPS biosynthesis to promote *Bacillus subtilis biofilm* formation. PLoS Genet. 2010;6(12):e1001243. Epub 2010/12/21. doi: 10.1371/journal.pgen.1001243. PubMed PMID: 21170308; PubMed Central PMCID: PMC3000366.

45. Wang R, Wang F, He R, Zhang R, Yuan J. The second messenger c-di-GMP adjusts motility and promotes surface aggregation of bacteria. Biophys J. 2018;115:2242–9.

46. Huangyutitham V, Guvener ZT, Harwood CS. Subcellular clustering of the phosphorylated WspR response regulator protein stimulates its diguanylate cyclase activity. mBio. 2013;4(3):e00242–13. Epub 2013/05/09. doi: 10.1128/mBio.00242-13. PubMed PMID: 23653447; PubMed Central PMCID: PMC3663191.

47. Bennett RR, Lee CK, De Anda J, Nealson KH, Yildiz FH, O’Toole GA, et al. Species-dependent hydrodynamics of flagellum-tethered bacteria in early biofilm development. J R Soc Interface. 2016;13(115):20150966. doi: 10.1098/rsif.2015.0966. PubMed PMID: 26864892; PubMed Central PMCID: PMCPMC4780565.

48. Luo Y, Zhao K, Baker AE, Kuchma SL, Coggan KA, Wolfgang MC, et al. A hierarchical cascade of second messengers regulates *Pseudomonas aeruginosa* surface behaviors. mBio. 2015;6(1):e02456–14. Epub 2015/01/30. doi: 10.1128/mBio.02456-14. PubMed PMID: 25626906; PubMed Central PMCID: PMC4324313.

49. Persat A, Inclan YF, Engel JN, Stone HA, Gitai Z. Type IV pili mechanochemically regulate virulence factors in *Pseudomonas aeruginosa*. Proc Natl Acad Sci U S A. 2015. Epub 2015/06/05. doi: 10.1073/pnas.1502025112. PubMed PMID: 26041805.

50. Lee CK, de Anda J, Baker AE, Bennett RR, Luo Y, Lee EY, et al. Multigenerational memory and adaptive adhesion in early bacterial biofilm communities. Proc Natl Acad Sci U S A. 2018;115(17):4471–6. Epub 2018/03/22. doi: 10.1073/pnas.1720071115. PubMed PMID: 29559526; PubMed Central PMCID: PMCPMC5924909.

51. Vogel HJ, Bonner DM. Acetylornithinase of *Escherichia coli:* partial purification and some properties. J Biol Chem. 1956;218:97–106.

52. O’Toole GA, Pratt LA, Watnick PI, Newman DK, Weaver VB, Kolter R. Genetic approaches to study of biofilms. Methods Enzymol. 1999;310:91–109.

53. Jain R, Kazmierczak BI. A Conservative Amino Acid Mutation in the Master Regulator FleQ Renders *Pseudomonas aeruginosa* Aflagellate. PLoS One. 2014;9(5):e97439. Epub 2014/05/16. doi: 10.1371/journal.pone.0097439. PubMed PMID: 24827992.

54. Giardine B, Riemer C, Hardison RC, Burhans R, Elnitski L, Shah P, et al. Galaxy: a platform for interactive large-scale genome analysis. Genome Res. 2005;15(10):1451–5. Epub 2005/09/20. doi: 10.1101/gr.4086505. PubMed PMID: 16169926; PubMed Central PMCID: PMCPMC1240089.

55. Li H, Durbin R. Fast and accurate short read alignment with Burrows-Wheeler transform. Bioinformatics. 2009;25(14):1754–60. Epub 2009/05/20. doi: 10.1093/bioinformatics/btp324. PubMed PMID: 19451168; PubMed Central PMCID: PMCPMC2705234.

56. Li H, Durbin R. Fast and accurate long-read alignment with Burrows-Wheeler transform. Bioinformatics. 2010;26(5):589–95. Epub 2010/01/19. doi: 10.1093/bioinformatics/btp698. PubMed PMID: 20080505; PubMed Central PMCID: PMCPMC2828108.

57. Langmead B, Trapnell C, Pop M, Salzberg SL. Ultrafast and memory-efficient alignment of short DNA sequences to the human genome. Genome Biol. 2009;10(3):R25. Epub 2009/03/06. doi: 10.1186/gb-2009-10-3-r25. PubMed PMID: 19261174; PubMed Central PMCID: PMCPMC2690996.

58. Li H, Handsaker B, Wysoker A, Fennell T, Ruan J, Homer N, et al. The Sequence Alignment/Map format and SAMtools. Bioinformatics. 2009;25(16):2078–9. Epub 2009/06/10. doi: 10.1093/bioinformatics/btp352. PubMed PMID: 19505943; PubMed Central PMCID: PMCPMC2723002.

59. Garrity-Ryan L, Kazmierczak B, Kowal R, Comolli J, Hauser A, Engel J. The arginine finger domain of ExoT is required for actin cytoskeleton disruption and inhibition of internalization of *Pseudomonas aeruginosa* by epithelial cells and macrophages. Infect Immun. 2000;68:7100–13.

60. de Kerchove AJ, Elimelech M. Impact of alginate conditioning film on deposition kinetics of motile and nonmotile *Pseudomonas aeruginosa* strains. Appl Environ Microbiol. 2007;73:5227–34.

61. Vallet-Gely I, Donovan KE, Fang R, Joung JK, Dove SL. Repression of phase-variable cup gene expression by H-NS-like proteins in *Pseudomonas aeruginosa*. Proc Natl Acad Sci U S A. 2005;102(31): 11082–7. Epub 2005/07/27. doi: 10.1073/pnas.0502663102. PubMed PMID: 16043713; PubMed Central PMCID: PMC1182424.

62. Dove SL, Hochschild A. Conversion of the omega subunit of *Escherichia coli* RNA polymerase into a transcriptional activator or an activation target. Genes Dev. 1998; 12:745–54.

63. Dufour YS, Gillet S, Frankel NW, Weibel DB, Emonet T. Direct Correlation between Motile Behavior and Protein Abundance in Single Cells. PLoS Comput Biol. 2016;12(9):e1005041. doi: 10.1371/journal.pcbi.1005041. PubMed PMID: 27599206.

64. Parthasarathy R. Rapid, accurate particle tracking by calculation of radial symmetry centers. Nat Methods. 2012;9(7):724–6. Epub 2012/06/13. doi: 10.1038/nmeth.2071. PubMed PMID: 22688415.

65. Jaqaman K, Loerke D, Mettlen M, Kuwata H, Grinstein S, Schmid SL, et al. Robust single-particle tracking in live-cell time-lapse sequences. Nat Methods. 2008;5(8):695–702. Epub 2008/07/22. doi: 10.1038/nmeth.1237. PubMed PMID: 18641657; PubMed Central PMCID: PMCPMC2747604.

66. Theves M, Taktikos J, Zaburdaev V, Stark H, Beta C. A bacterial swimmer with two alternating speeds of propagation. Biophys J. 2013;105(8):1915–24. doi: 10.1016/j.bpj.2013.08.047. PubMed PMID: 24138867; PubMed Central PMCID: PMCPMC3797586.

