## Supplemental Table 1 for "Modulation of Flagellar Rotation in Surface-Attached Bacteria: A Circuit for Rapid Surface-Sensing"

**Supplemental Table 1. Mutations mapped in FlhF(R251G) suppressors**

| **Suppressor** | **Mutations mapped by sequencing ***  *(numbers indicate last correct aa in indel and nonsense mutations)* | |
| --- | --- | --- |
|  | **outside *vfr*** | **in *vfr*** |
| **Mutations mapped by whole genome sequencing** | | |
| EMS 15 | *PA0916(G163S), wrbA(G197E), purT(A85T), PA3969(stop@aa133)* | *vfr(stop157)* |
| EMS 63 | *PA0544(R224Q), alkA(G182D), pslM(T28A), bexR(T229A), katB(G428D), secG(A102T), PA5021(V524M)* | *vfr(C183Y)* |
| UV 07 | - | *Δvfr* |
| UV 37 | - | *vfr(indel32) [-1bp]* |
| WO 11 | - | *vfr(indel32) [+1bp]* |
| UV 24 | - | *vfr(indel147) [-39bp]* |
| UV 16 | - | *vfr(indel147) [+1bp]* |
| NG 01 | - | *vfr(G73D)* |
| WO 04 | - | *vfr(L75Q)* |
| NG 36 | - | *vfr(G76E)* |
| NG 38 | - | *vfr(G122D)* |
| UV 33 | - | *vfr(L155P)* |
| NG 11 | *PA0134(S201N), PA5370(G333D)* | *vfr(G137D)* |
| WO 35 | *PA5318(F145L),* ***fimV(L7P)*** | - |
| NG 04 | ***fimV(L7P)*** | - |
| **Mutations mapped by targeted gene sequencing** | | |
| UV 23 | ND | *vfr(Y25D)* |
| NG 18 | ND | *vfr(G47S)* |
| UV 10 | ND | *vfr(Y65D)* |
| NG 13 | ND | *vfr(A126V)* |
| WO 23 | ND | *vfr(R174L)* |
| NG 17 | ND | *vfr(G182D)* |
| NG 08 | ND | *vfr(G189E)* |
| NG 12 | ND | *vfr(G205R)* |
| NG 25 | ***fimV(stop115)*** | - |
| NG 30 | ***fimV(stop826)*** | - |
