## Supplemental Table 2 for "Modulation of Flagellar Rotation in Surface-Attached Bacteria: A Circuit for Rapid Surface-Sensing"

**S1 Table. Bacterial strains and plasmids used in this study**

| **Strain or plasmid** | **Description or relevant genotype** | **Source or reference** |
| --- | --- | --- |
| *E. coli* strains |  |  |
| DH5α | Used for subcloning | Invitrogen |
| XL1-Blue | Used for subcloning | Stratagene |
| S17-1 | Used for mating constructs into *P. aeruginosa* | [1] |
| KDZif1∆Z | Used for bacterial two-hybrid assay | [2] |
| *P. aeruginosa* strains |  |  |
| PA14 | Wild type isolate | F. Ausubel |
| PAK | Wild type isolate | J. Mattick |
| PAK *∆fliC* | Deletion of *fliC (PA1092)* | J. Mattick |
| PAK *∆pilA* | Deletion of *pilA (PA4525)* | [3] |
| PAK *∆flhF* | In-frame deletion of amino acids 7-430 of FlhF *(PA1453)* | [4] |
| PAK *∆flhF pilA* | Deletion of *flhF* and *pilA* | This work |
| PAK *∆motAB* | Deletion of *motA (PA4954)* and *motB (PA4953)* | This work |
| PAK *∆motCD* | Deletion of *motC (PA1460)* and *motD (PA1461)* | This work |
| PAK *∆flhF ∆motAB* | Deletion of *motA (PA4954)* and *motB (PA4953)* in *∆flhF* | This work |
| PAK *∆flhF ∆motCD* | Deletion of *motC (PA1460)* and *motD (PA1461)* in *∆flhF* | This work |
| PAK *∆cheR1* | Deletion of *cheR1 (PA3348)* | This work |
| PAK *∆cheZ* | Deletion of *cheZ (PA1457)* | This work |
| PAK *∆cbpA* | Deletion of *cbpA (PA4704)* | This work |
| PAK *attB::flhF* | *FlhF-His6* under the control of an arabinose promoter integrated at the *attB* site (*attB*::p_ARA_ *flhF-His6)* | This work |
| PAK *attB::flhF(R251G)* | *FlhF(R251G)-His6* under the control of an arabinose promoter integrated at the *attB* site (*attB*::p_ARA_ *flhF (R251G)-His6)* | This work |
| PAK *∆flhF attB::flhF(R251G)* | *FlhF(R251G)-His6* under the control of an arabinose promoter integrated at the *attB* site in *∆flhF* background  (*attB*::p_ARA_ *flhF (R251G)-His6)* | [3] |
| NG01 | Suppressor of PAK *∆flhF* + *attB::flhF(R251G)* with *vfr(G73D)* | This work |
| PA14 *fimV::Tn* | Transposon insertion disrupting *fimV* | G. O’Toole |
| PA14 *pilI::Tn* | Transposon insertion disrupting *pilI* | G. O’Toole |
| PA14 *pilJ::Tn* | Transposon insertion disrupting *pilJ* | G. O’Toole |
| PA14 *pilG::Tn* | Transposon insertion disrupting *pilG* | G. O’Toole |
| PA14 *pilQ::Tn* | Transposon insertion disrupting *pilQ* | G. O’Toole |
| PAK *fimV(L7P)* | *fimV(L7P)* under its native promoter at the *fimV* site | This work |
| PAK *fimV(ΔC)* | *fimV(1-769)* under its native promoter at the *fimV* site | This work |
| PAK *fimV-BB2* | *fimV-BB2* under its native promoter at the *fimV* site | This work |
| PAK *fimV(L7P)-BB2* | *fimV(L7P)-BB2* under its native promoter at the *fimV* site | This work |
| Plasmids |  |  |
| pUCP | Expression vector; Ap^R^ | [5] |
| pFlhF(R251G) | *flhF(R251G)* cloned in pUCP under *lac* promoter; Ap^R^ | This work |
| pMotAB | *motA* and *motB* cloned in pUCP under *lac* promoter; Ap^R^ | This work |
| pMotCD | *motC* and *motD* cloned in pUCP under *lac* promoter; Ap^R^ | This work |
| pFleQ | *fleQ* (*PA1097*) cloned in pUCP under *lac* promoter; Ap^R^ | [6] |
| pDONRX | Gateway-adapted suicide vector; Gm^R^ | [7] |
| pDONRX/*fimV-BB2* | Shuttle vector to introduce *fimV-BB2* into chromosome; Gm^R^ | This work |
| pDONRX/*fimV(L7P)* | Shuttle vector to introduce *fimV(L7P)* into chromosome; Gm^R^ | This work |
| pDONRX/*fimV(ΔC)* | Shuttle vector to introduce *fimV(1-769)* into chromosome; Gm^R^ | This work |
| pDONRX*/ΔmotAB* | Shuttle vector to delete *motA (PA4954)* and *motB (PA4953)*; Gm^R^ | This work |
| pDONRX/*ΔmotCD* | Shuttle vector to delete *motC (PA1460)* and *motD (PA1461)*; Gm^R^ | This work |
| pDONRX/*ΔcheR1* | Shuttle vector to delete *cheR1 (PA3348)*; Gm^R^ | This work |
| pDONRX/*ΔcheZ* | Shuttle vector to delete *cheZ (PA1457)*; Gm^R^ | This work |
| pDONRX/*ΔcbpA* | Shuttle vector to delete *cbpA (PA4704)*; Gm^R^ | This work |
| pEX18Gm/*ΔpilA* | Shuttle vector to delete *pilA* (*PA4525*) ; Gm^R^ Tc^R^ | [8] |
| pMMB67EH | Shuttle vector; Ap^R^ | [9] |
| pMMB67EH/Vfr | Shuttle vector to express Vfr; Ap^R^ | [10] |
| pBRGPω | Vector for bacterial two hybrid assay with a C-TD reporter fusion of ω subunit of *E. coli* RNA polymerase; Ap^R^ | [2] |
| pACTR-AP-Zif | Vector for bacterial two hybrid assay with a C-TD reporter fusion of zinc finger DNA-binding domain of Zif; Tc^R^ | [2] |
| pBRGP / *flhF(WT)-ω* | Vector for bacterial two hybrid with FlhF(WT)-ω fusion; Ap^R^ | [3] |
| pBRGP / *flhF(R251G)-ω* | Vector for bacterial two hybrid with FlhF(R251G)-ω fusion; Ap^R^ | [3] |
| pBRGP / *flhF(K222A)-ω* | Vector for bacterial two hybrid with FlhF(K222A)-ω fusion; Ap^R^ | [3] |
| pBRGP / *flhF(D294A)-ω* | Vector for bacterial two hybrid with FlhF(D294A)-ω fusion; Ap^R^ | [3] |
| pACTR / *flhF(WT)*-Zif | Vector for bacterial two hybrid with FlhF(WT)-Zif fusion; Tc^R^ | [3] |
| pACTR / *flhF(R251G)*-Zif | Vector for bacterial two hybrid with FlhF(R251G)-Zif fusion; Tc^R^ | [3] |
| pACTR / *flhF(K222A)*-Zif | Vector for bacterial two hybrid with FlhF(K222A)-Zif fusion; Tc^R^ | [3] |
| pACTR / *flhF(D294A)*-Zif | Vector for bacterial two hybrid with FlhF(D294A)-Zif fusion; Tc^R^ | [3] |
| pBRGP / *flhF(L298R,P299L)-ω* | Vector for bacterial two hybrid with FlhF(L298R,P299L)-ω fusion; Ap^R^ | This work |
| pACTR / *flhF(*L298R,P299L)-Zif | Vector for bacterial two hybrid with FlhF(L298R,P299L)-Zif fusion; Tc^R^ | This work |
| pBRGP / *fimV(cyto)-ω* | Vector for bacterial two hybrid with FimV(aa 491-919)-ω fusion; Ap^R^ | This work |
| pBRGP / *fimV(cytoΔC)-ω* | Vector for bacterial two hybrid with FimV(aa 491-769)-ω fusion (aa 491-769); Ap^R^ | This work |
| pACTR / *fimV(cyto)*-Zif | Vector for bacterial two hybrid with FimV(aa 491-919)-Zif fusion; Tc^R^ | This work |
| pACTR / *fimV(cytoΔC)*-Zif | Vector for bacterial two hybrid with FimV(aa 491-769)-Zif fusion (aa 491-769); Tc^R^ | This work |
| pBRGP / *vfr-ω* | Vector for bacterial two hybrid with Vfr-ω fusion; Ap^R^ | This work |
| pACTR / *vfr*-Zif | Vector for bacterial two hybrid with Vfr-Zif fusion; Tc^R^ | This work |
| pBRGP / *fliF-ω* | Vector for bacterial two hybrid with FliF-ω fusion; Ap^R^ | This work |
| pACTR / *fliF*-Zif | Vector for bacterial two hybrid with FliF-Zif fusion; Tc^R^ | This work |
| pBRGP / *fliG-ω* | Vector for bacterial two hybrid with FliG-ω fusion; Ap^R^ | This work |
| pACTR / *fliG*-Zif | Vector for bacterial two hybrid with FliG-Zif fusion; Tc^R^ | This work |
| pBRGP / *FliM-ω* | Vector for bacterial two hybrid with FliM-ω fusion; Ap^R^ | This work |
| pACTR / *FliM*-Zif | Vector for bacterial two hybrid with FliM-Zif fusion; Tc^R^ | This work |
| pBRGP / *FliN-ω* | Vector for bacterial two hybrid with FliN-ω fusion; Ap^R^ | This work |
| pACTR / *FliN*-Zif | Vector for bacterial two hybrid with FliN-Zif fusion; Tc^R^ | This work |
| pBRGP / *motA-ω* | Vector for bacterial two hybrid with MotA-ω fusion; Ap^R^ | This work |
| pACTR / *motA*-Zif | Vector for bacterial two hybrid with MotA-Zif fusion; Tc^R^ | This work |
| pBRGP / *motB-ω* | Vector for bacterial two hybrid with MotB-ω fusion; Ap^R^ | This work |
| pACTR / *motB*-Zif | Vector for bacterial two hybrid with MotB-Zif fusion; Tc^R^ | This work |
| pBRGP / *motC-ω* | Vector for bacterial two hybrid with MotC-ω fusion; Ap^R^ | This work |
| pACTR / *motC*-Zif | Vector for bacterial two hybrid with MotC-Zif fusion; Tc^R^ | This work |
| pBRGP / *motD-ω* | Vector for bacterial two hybrid with MotD-ω fusion; Ap^R^ | This work |
| pACTR / *motD*-Zif | Vector for bacterial two hybrid with MotD-Zif fusion; Tc^R^ | This work |

**References**

1.	Simon R, Priefer U, Puhler A. A broad host range mobilization system for in vivo genetic engineering: transposon mutagenesis in Gram negative bacteria. Biotechnology. 1983;1:784-91.

2.	Vallet-Gely I, Donovan KE, Fang R, Joung JK, Dove SL. Repression of phase-variable cup gene expression by H-NS-like proteins in *Pseudomonas aeruginosa*. Proc Natl Acad Sci U S A. 2005;102(31):11082-7.

3.	Schniederberend M, Abdurachim K, Murray TS, Kazmierczak BI. The GTPase Activity of FlhF Is Dispensable for Flagellar Localization, but Not Motility, in *Pseudomonas aeruginosa*. J Bacteriol. 2013;195(5):1051-60.

4.	Murray TS, Kazmierczak BI. FlhF is required for swimming and swarming in *Pseudomonas aeruginosa*. J Bacteriol. 2006;188(19):6995-7004.

5.	Watson AA, Alm RA, Mattick JS. Construction of improved vectors for protein production in *Pseudomonas aeruginosa*. Gene. 1996;172:163-4.

6.	Jain R, Kazmierczak BI. A Conservative Amino Acid Mutation in the Master Regulator FleQ Renders *Pseudomonas aeruginosa* Aflagellate. PLoS One. 2014;9(5):e97439.

7.	Fulcher NB, Holliday PM, Klem E, Cann MJ, Wolfgang MC. The *Pseudomonas aeruginosa* Chp chemosensory system regulates intracellular cAMP levels by modulating adenylate cyclase activity. Mol Microbiol. 2010;76(4):889-904.

8.	de Kerchove AJ, Elimelech M. Impact of alginate conditioning film on deposition kinetics of motile and nonmotile *Pseudomonas* *aeruginosa* strains. Appl Environ Microbiol. 2007;73:5227-34.

9.	Furste JP, Pansegrau W, Frank R, Blocker H, Scholz P, Bagdasarian M, et al. Molecular cloning of the plasmid RP4 primase region in a multi-host range *tacP* expression vector. Gene. 1986;48:119-31.

10.	Wolfgang MC, Lee VT, Gilmore ME, Lory S. Coordinate regulation of bacterial genes by a novel adenylate cyclase signaling pathway. Devlop Cell. 2003;4:253-63.
