## Supplemental Table 3 for "Modulation of Flagellar Rotation in Surface-Attached Bacteria: A Circuit for Rapid Surface-Sensing"

**S2 Table. Primers used in this study**

| Purpose | Primer | Sequence (5’ - 3’) |
| --- | --- | --- |
| **Chromosomal constructs** | | |
| *fimV(L7P)* | attB1-FimV-forw | GGGG ACA AGT TTG TAC AAA AAA GCA GGC T CTG CGC CTG CAC CTG GGA |
|  | attB2-FimV-rev | GGGG AC CAC TTT GTA CAA GAA AGC TGG GT C CTG CCT CTC CGT CTC CAG |
| *fimV-BB2*  or  (*fimV(L7P)-BB2*) | attB1-FimV2-forw | GGGG ACA AGT TTG TAC AAA AAA GCA GGC T GAC GAC CTG GGC GGC GAC |
|  | BB2-FimV2-rev | GTC GAG CGG GTC CTG GTT GGT GTG CAC CTC GGC CAG GCG CTC CAG CAA CTC |
|  | BB2-FimV2-for | GAG GTG CAC ACC AAC CAG GAC CCG CTC GAC TGA TCC GGA AAT GAA GCG ACC CTC |
|  | attB2-FimV2-rev | GGGG AC CAC TTT GTA CAA GAA AGC TGG GT CAG ATA GAG TCC GTA GGG GTG |
| *fimV(ΔC)* | attB1-FimVCT-forw | GGGG ACA AGT TTG TAC AAA AAA GCA GGC T AGG TCG AGA AGA CCA CCG CGC |
|  | FimVCT-rev | GGT CGC TTC ATT TCC GGA TCA GGC GGG CAC TTC GTC GTT CAG |
|  | FimVCT-forw | CTG AAC GAC GAA GTG CCC GCC TGA TCC GGA AAT GAA GCG ACC |
|  | attB2-FimVCT-rev | GGGG AC CAC TTT GTA CAA GAA AGC TGG GT ATT CGA TCG GCC GCT CGC CGG |
| *ΔmotAB* | attB1-MotAB-KO-forw | GGGG ACA AGT TTG TAC AAA AAA GCA GGC T GCC AGG CAG ATC GGC AGC AGG |
|  | MotAB-KO-rev | TGA ACC **TCA** GTC GCC CTT GAT GAT GAT TTT TGA **CAT** GAG GAC |
|  | MotAB-KO-forw | GTC CTC **ATG** TCA AAA ATC ATC ATC AAG GGC GAC **TGA** GGT TCA |
|  | attB2-MotAB-KO-rev | GGGG AC CAC TTT GTA CAA GAA AGC TGG GT CCC GGA CAT GCG CGG GCT GCT |
| *ΔmotCD* | attB1-MotCD-KO-forw | GGGG ACA AGT TTG TAC AAA AAA GCA GGC T GCC GGC GCG TCG CCG GCC CCG |
|  | MotCD-KO-rev | GCG CGC **TCA** TGG CGA AGG CGA GCT GAG CAC ATC **CAT** CAG CGC |
|  | MotCD-KO-forw | GCG CTG **ATG** GAT GTG CTC AGC TCG CCT TCG CCA **TGA** GCG CGC |
|  | attB2-MotCD-KO-rev | GGGG AC CAC TTT GTA CAA GAA AGC TGG GT AAT CCG CAG CGT GCT CAG CGA |
| *ΔcheR1* | attB1-CheR1-KO-forw | GGGG ACA AGT TTG TAC AAA AAA GCA GGC T AGG CAC CAT TCC GAT CCT CGA |
|  | CheR1-KO-rev | CTT GCG **CTA** CTT GGC CCG GTA ATT AGC TGC CGA **CAC** GCA TAA |
|  | CheR1-KO-forw | TTA TGC **GTG** TCG GCA GCT AAT TAC CGG GCC AAG **TAG** CGC AAG |
|  | attB2-CheR1-KO-rev | GGGG AC CAC TTT GTA CAA GAA AGC TGG GT GCA TCA TCA CGT CCA TCA GGA |
| *ΔcheZ* | attB1-CheZ-KO-forw | GGGG ACA AGT TTG TAC AAA AAA GCA GGC T TGT ACA GCT TCG ACG ACC TGT |
|  | CheZ-KO-rev | GGC GGG **TCA** GAA ACC CAG GCT GTT ACC AAG CAC **CAT** GAC ACC |
|  | CheZ-KO-forw | GGT GTC **ATG** GTG CTT GGT AAC AGC CTG GGT TTC **TGA** CCC GCC |
|  | attB2-CheZ-KO-rev | GGGG AC CAC TTT GTA CAA GAA AGC TGG GT A ACT CGG CGT CGC TGA TCT C |
| *ΔcbpA* | attB1-CbpA-KO-forw | GGGG ACA AGT TTG TAC AAA AAA GCA GGC T AGA TAT CCG GTA CGT ATT TCC |
|  | CbpA-KO-rev | GC TGT **TCA** GGC GCT GCG GCG GCC GAG TAG ATA **CAT** GCC CGA |
|  | CbpA-KO-forw | TCG GGC **ATG** TAT CTA CTC GGC CGC CGC AGC GCC **TGA** ACA GC |
|  | attB2-CbpA-KO-rev | GGGG AC CAC TTT GTA CAA GAA AGC TGG GT GCC GAG CGT CTT CGC GAT GCG |
| **Bacterial two hybrid constructs** | | |
| *fimV(cyto)* | NdeI-FimV-forw | **CAT ATG** AAA GAG AAG GAA GAA GCC CAG GCT TTC GCC GCG |
|  | NotI-FimV-rev | **GC GGC CGC** GGC CAG GCG CTC CAG CAA CTC GCG GGC TTC CGC |
| *fimV(ctyoΔC)* | NotI-FimV2-rev | **GC GGC CGC** GTT GTC GGC GGG CGC CGC |
| *vfr* | NdeI-Vfr-forw | **CAT ATG** GTA GCT ATT ACC CAC ACA CCC AAA CTC AAA CAC |
|  | NotI-Vfr-rev | **GC GGC CGC** GCG GGT GCC GAA GAC CAC CAT GGT CTT TCC TTT |
| *fliF* | NdeI-FliF-forw | TATAGA **CAT ATG** GCC GAT GCA CTG ATC GAC AGC CAG GTT |
|  | NotI-FliF-rev | TATAGA **GCGGCCGC** CTC ATC GGC GTT GAT CCA CTC TTT CAC |
| *fliG* | NdeI-FliG-forw | TATAGA **CAT ATG** AGT GAG AAT CGT CTC GCC GCC AAA CTG |
|  | NotI-FliG-rev | TATAGA **GCGGCCGC** GAT CAT CTC CTC GCC ACC CTT GCC GCC CAG |
| *fliM* | NdeI-FliM-forw | TATAGA **CAT ATG** GCC GTG CAA GAT CTG CTT TCC CAG GAT |
|  | NotI-FliM-rev | TATAGA **GCGGCCGC** GCG CGA GCG CTC GAC CGC TTC GAG AAT |
| *fliN* | NdeI-FliN-forw | TATAGA **CAT ATG** GCA GAC GAA GAA AAA GTG ACC ACC GAG |
|  | NotI-FliN-rev | TATAGA **GCGGCCGC** GCG CAG CTT CTT GAT GCG TTC GCT GGG ACT |
| *motA* | NdeI-MotA-forw | TATAGA **CAT ATG** TCA AAA ATC ATC GGC ATC ATC GTC GTT |
|  | PspOMI-MotA-rev | TATAGA **GGGCCC** A GCG GCC GCG CAC GGC CTG CTC GAG CTC |
| *motB* | NdeI-MotB-forw | TATAGA **CAT ATG** GAC AAT AAC CAG CCG ATC ATC GTC AAG |
|  | NotI-MotB-rev | TATAGA **GCGGCCGC** GTC GCC CTT GAT CTG CTC CAG CTT CAG |
| *motC* | NdeI-MotC-forw | TATAGA **CAT ATG** GAT GTG CTC AGC CTG GTC GGG ATC ATC |
|  | NotI-MotC-rev | TATAGA **GCGGCCGC** GTC CAT GAA GCC TTG CAG CTT CAA CTC |
| *motD* | NdeI-MotD-forw | TATAGA **CAT ATG** CAG CGC CGC CGT CGC CAC CAG GAG GAA |
|  | PspOMI-MotD-rev | TATAGA **GGGCCC** A TGG CGA AGG CGA CGG GGA ATT GAC GGT |

Bold = start or stop codon; Underlined = *attB* site; Bold and underlined = restriction site
