## Supplementary figures and images for "Modulation of Flagellar Rotation in Surface-Attached Bacteria: A Circuit for Rapid Surface-Sensing"

### S1 Figure

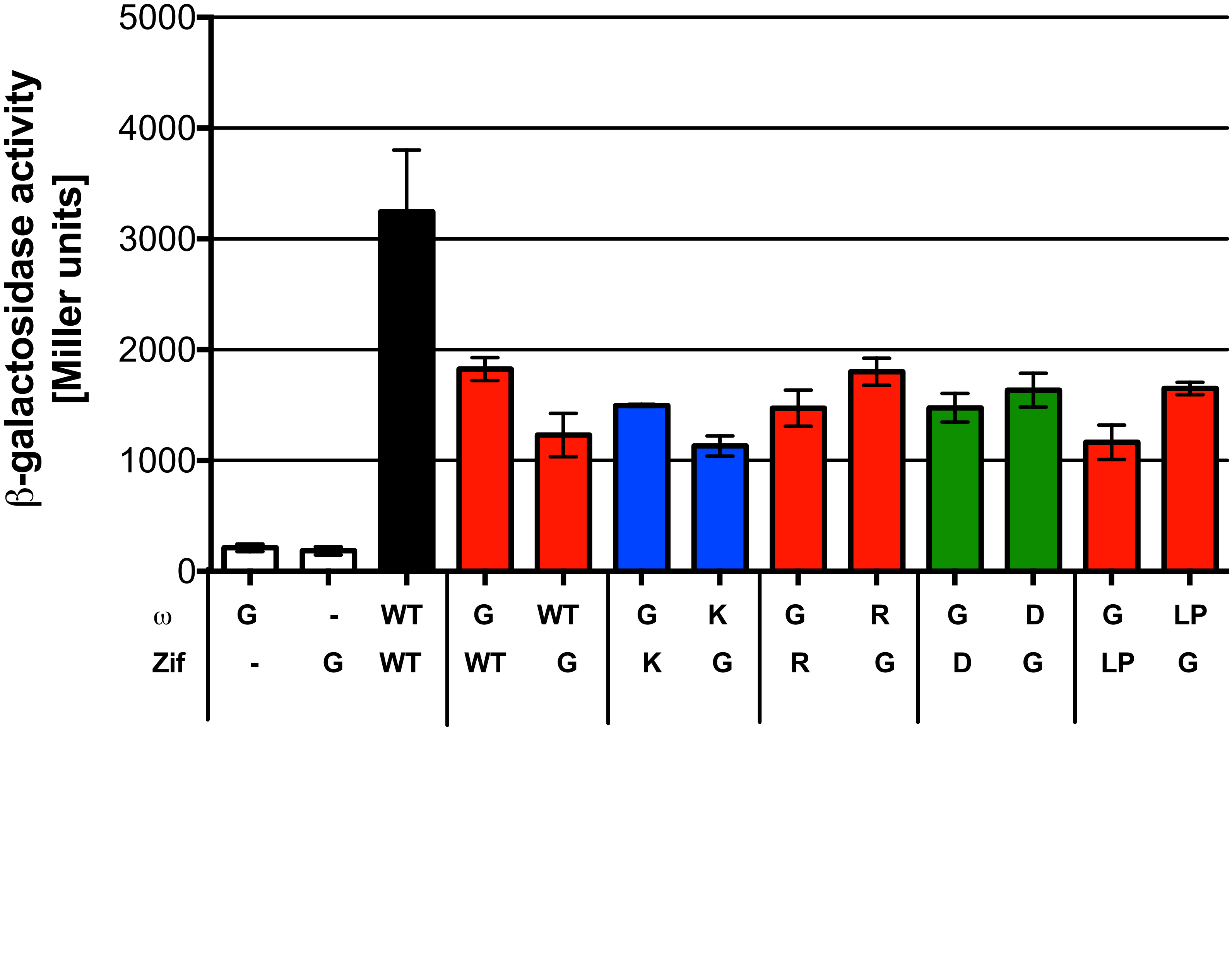

### S2 Figure

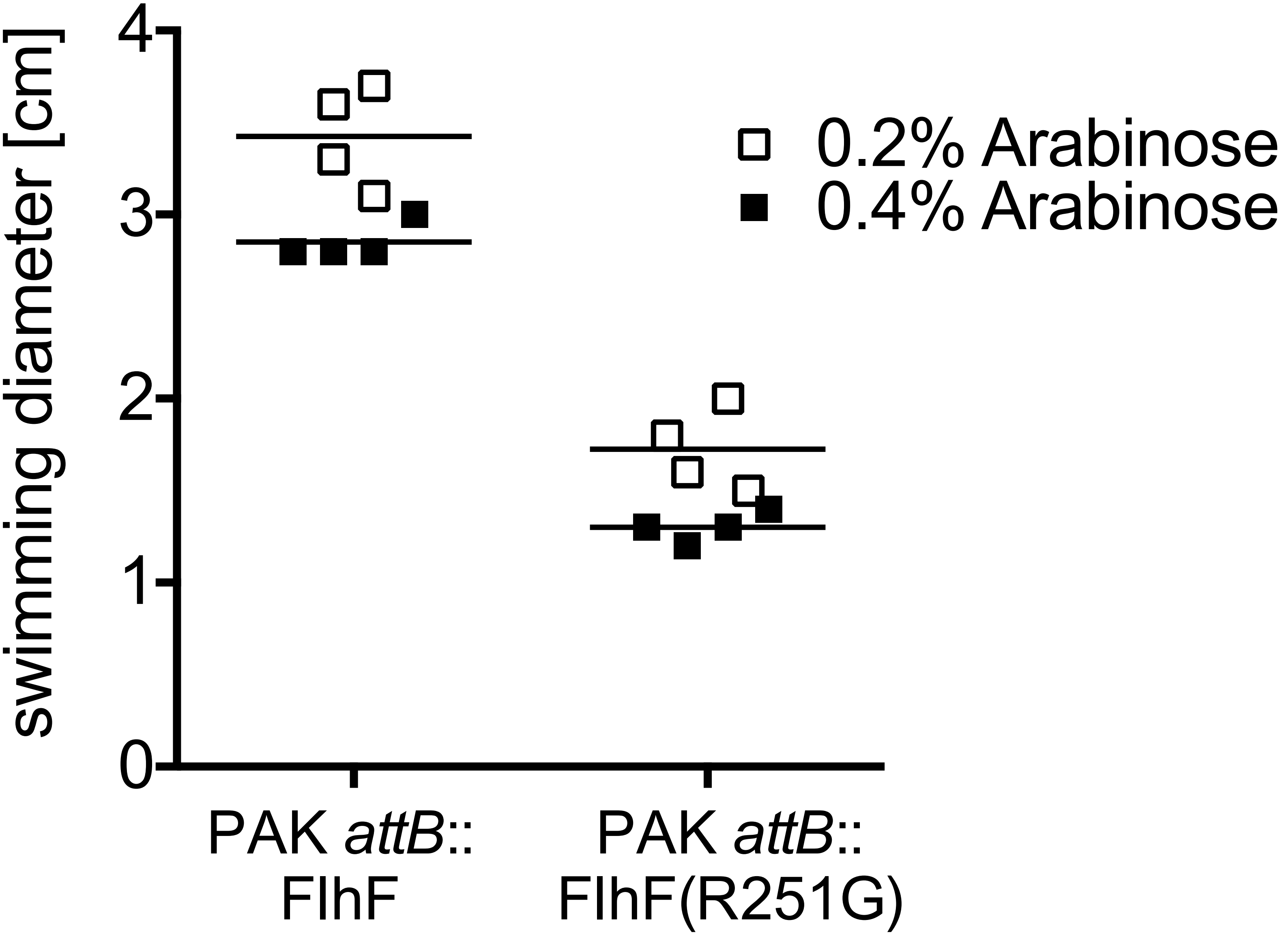

### S3 Figure

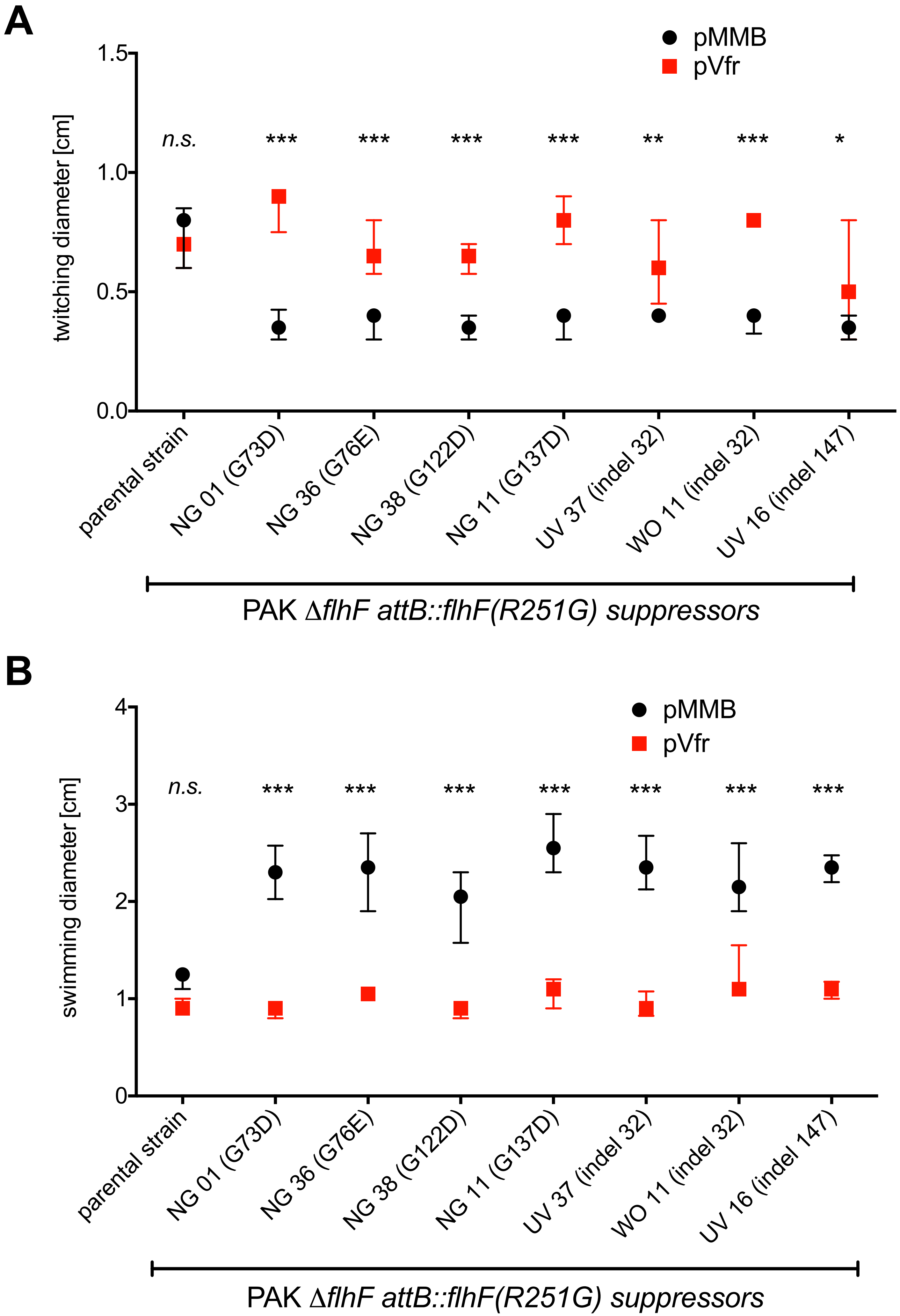

### S4 Figure

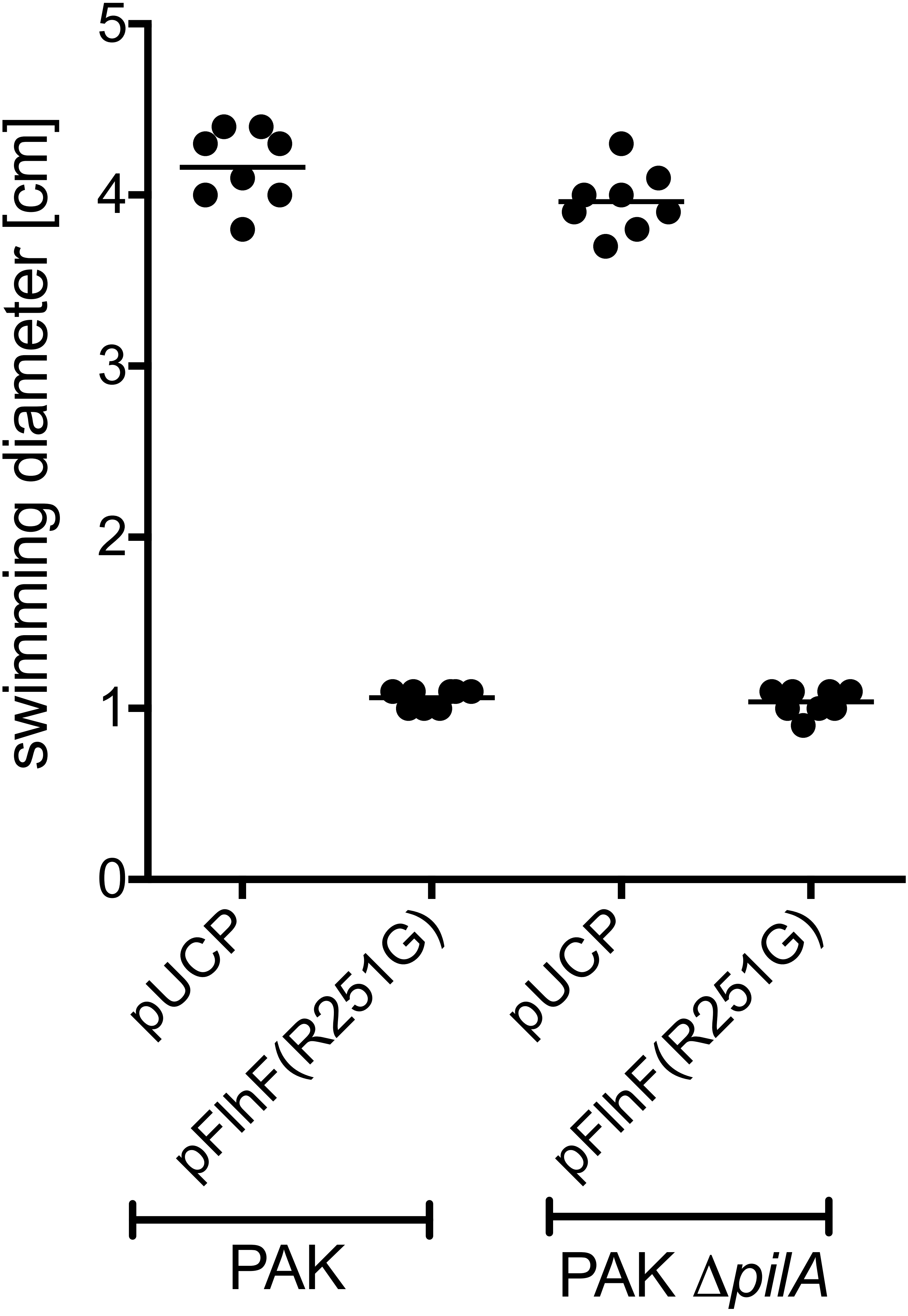

### S5 Figure

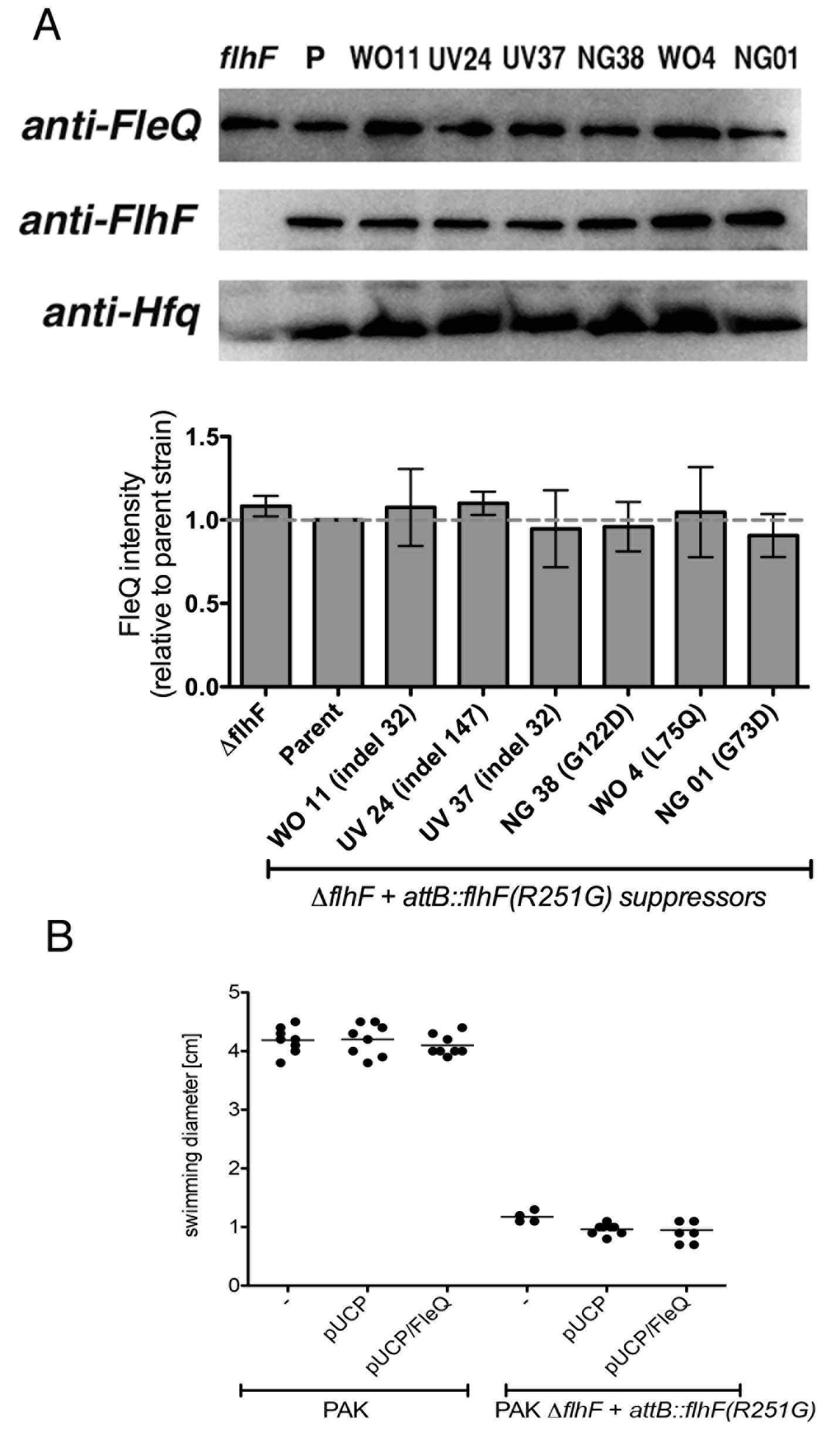

### S6 Figure

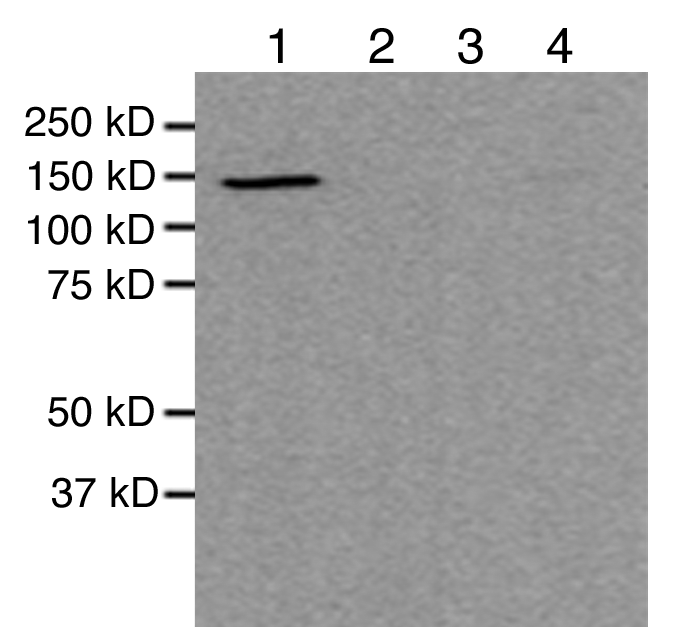

### S7 Figure

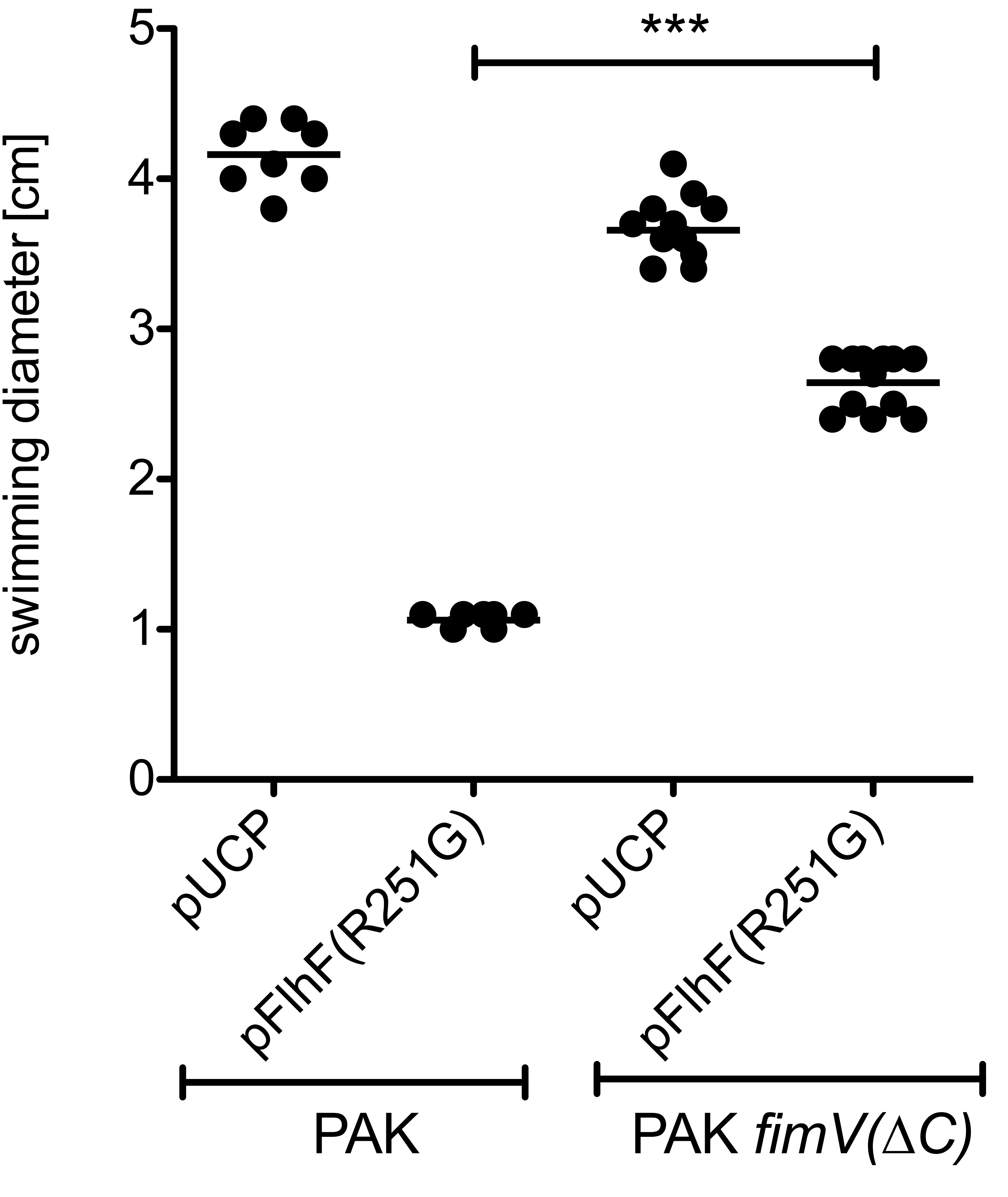

### S8 Figure

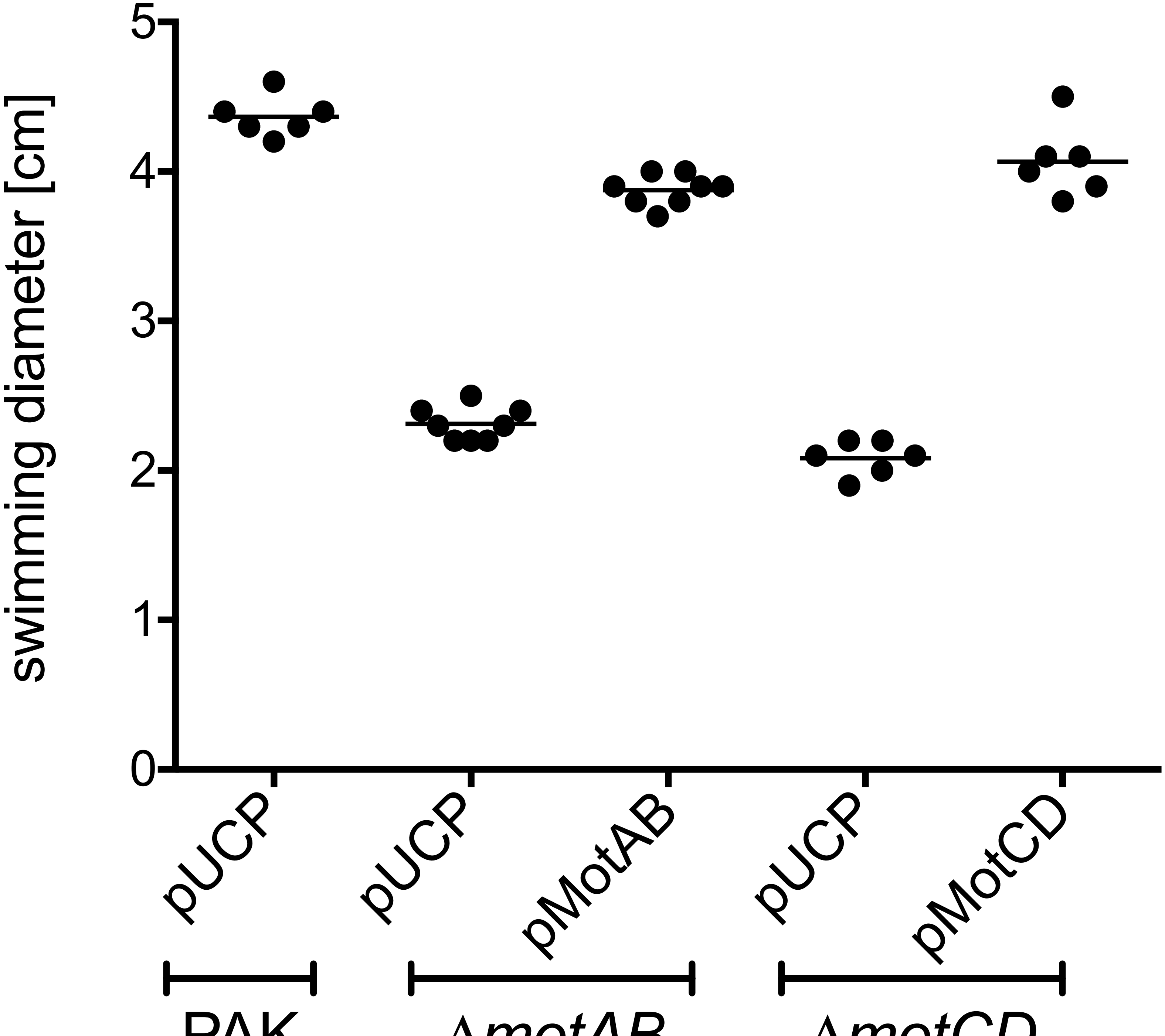

### S9 Figure

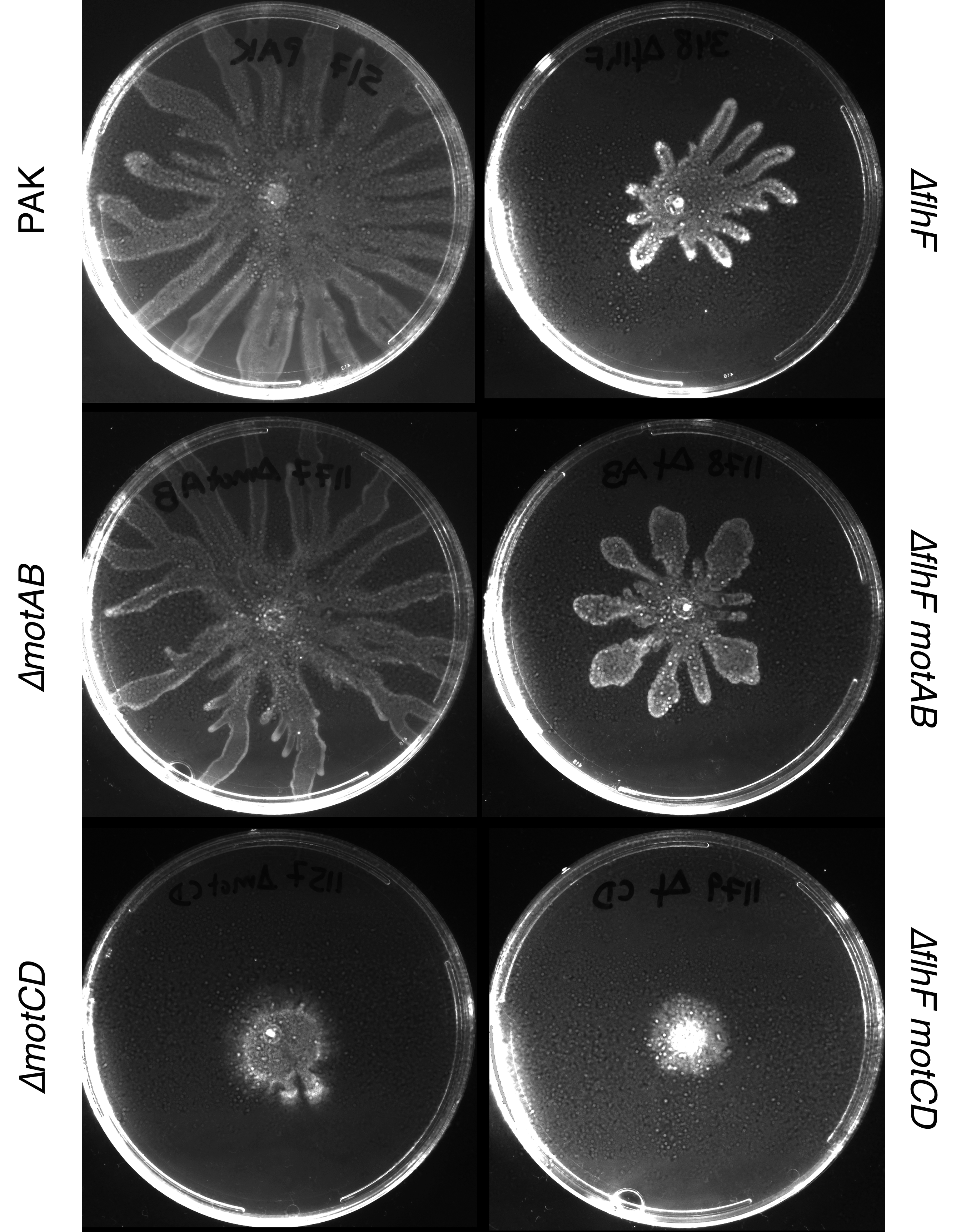

### S10 Figure

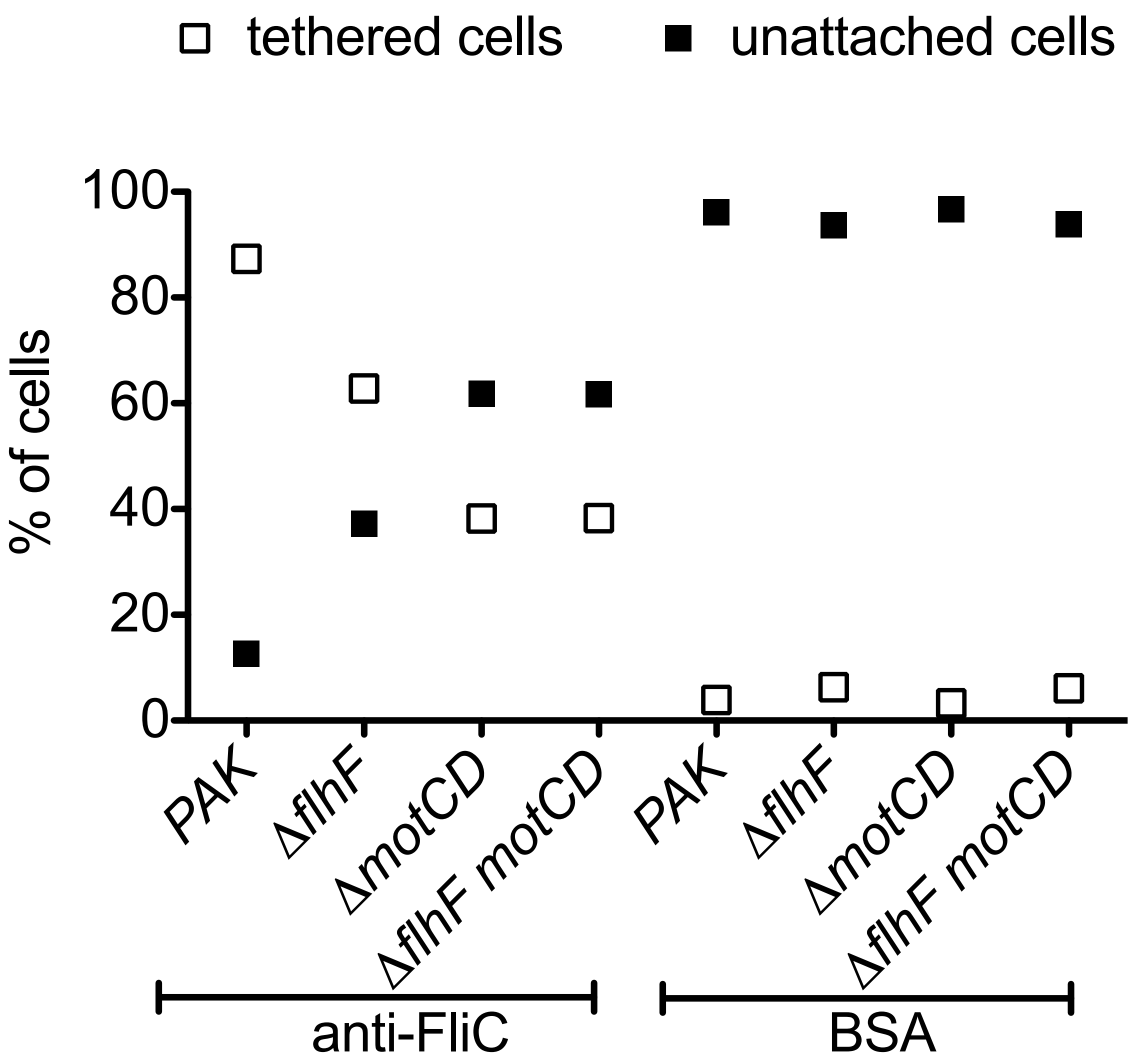


| | tethered | spinning (%) |
|---|---|---|
| PAK | 351 | 80 (23%) |
| *ΔflhF* | 268 | 107 (40%) |
| *ΔmotCD* | 131 | 5 (3.8%) |
| *ΔflhF motCD* | 200 | 15 (7.5%) |
